# Structural insights into autophosphorylation-mediated heat shock response by the protein-arginine kinase McsAB

**DOI:** 10.1101/2025.11.26.690658

**Authors:** Md Arifuzzaman, Kunwoong Park, Eunju Kwon, Dong Young Kim

## Abstract

McsAB is a protein-arginine kinase that activates heat shock genes by inhibiting the transcriptional repressor CtsR. However, the mechanism by which McsAB suppresses CtsR during heat shock remains unclear. Here, we present cryo-electron microscopy structures of *Bacillus subtilis* McsAZ2-McsB and its complex with CtsR. Our structural analyses reveal that McsAZ2-McsB forms a rod-shaped heterotetramer, with two of these heterotetramers assembling into an asymmetrically crossing shape through phosphorylated arginine residues (pArg194 and pArg333) of McsB. The CtsR dimer binds to the groove of the McsAZ2-McsB oligomer via pArg115 of McsA and pArg194 of McsB. Anisotropy experiments indicate that CtsR dissociates slowly from its operator during heat shock, and McsAB facilitates this process in an ATP-dependent manner. Furthermore, mutations in the arginine residues of McsAB reduce its ability to hijack CtsR during heat shock. These results demonstrate that autophosphorylation of McsAB is essential for a rapid and effective heat shock response.

## INTRODUCTION

Cells require specific temperature ranges for optimal growth. Upon exposure to temperatures beyond their optimal range, cellular defense systems are activated, enabling survival in altered conditions^1^. This response is primarily regulated by the transcription of heat shock genes^2,3^. Heat- responsive transcription factors induce the overexpression of these genes in response to heat shock. In *Bacillus subtilis*, transcription of heat shock genes is controlled by several factors, including transcriptional repressors (CtsR and HrcA), the heat shock sigma factor (SigB), and the two-component system CssS-CssR^4,5^. CtsR is a transcriptional repressor that binds to operator DNA located near the promoter^6–8^. The CtsR regulon, a set of genes regulated by CtsR, includes the *clpC* operon (comprising *ctsR*, *mcsA*, *mcsB*, and *clpC*), as well as *clpE* and *clpP*^6,7^. Under normal growth conditions, CtsR binds to the operator and represses transcription of the regulon. However, upon heat shock, CtsR dissociates, resulting in the activation of these genes^6–8^.

Two mechanisms have been proposed for CtsR inactivation during heat shock. The first suggests that CtsR acts as a heat sensor, dissociating from the operator upon exposure to elevated temperatures^9^. Electrophoretic mobility shift assays have shown that CtsR can dissociate from the operator at 50 °C^9^. The second mechanism involves the removal of CtsR from the operator through hijacking by McsAB^10–12^. *Bacillus subtilis* mutants lacking either *mcsA* or *mcsB* exhibit reduced transcription of the CtsR regulon under heat shock conditions^11,13^.

McsAB is a protein-arginine kinase that phosphorylates arginine residues in target proteins^10^. In this complex, McsB functions as the kinase subunit, and McsA acts as a regulatory subunit that activates McsB kinase activity^11,14,15^. McsA binds to McsB through the second zinc-coordinating motif and a subsequent loop (Z2)^15,16^. This interaction enhances the integrity of the active site and mediates the assembly of the McsAB oligomer^13,15^. Hundreds of arginine-specific phosphorylation sites have been identified in *B. subtilis* and *Staphylococcus aureus*^14,17–20^. Proteins tagged with phosphorylated arginine (pArg) by McsAB kinase are targeted to degradation by the ClpCP protease, indicating that pArg functions as a degradation signal in gram-positive bacteria, similar to the role of ubiquitin in eukaryotes^8,14^. Notably, the heat shock repressors CtsR and HrcA are primary targets for ClpCP-mediated degradation through arginine phosphorylation under heat shock conditions^4,10,14^.

McsAB kinase phosphorylates arginine residues in CtsR, leading to the degradation of pArg- tagged CtsR by the ClpCP protease^10,14^. Additionally, McsAB restricts the operator binding of CtsR through phosphorylation. CtsR loses its affinity for the operator when the residue R61, located on the operator-binding surface, is phosphorylated^10,11,21^. These suggest that the inactivation of CtsR during heat shock may be due to its phosphorylation by McsAB. However, pArg-independent degradation of CtsR by ClpCP has also been observed^9^. Even when all arginine residues in CtsR are replaced with lysine, ClpCP continues to degrade CtsR, suggesting that CtsR may be inactivated through a mechanism other than CtsR-phosphorylation^9^. Consequently, the specific mechanism by which McsAB regulates CtsR-dependent transcription under heat shock conditions remains unclear.

To gain a comprehensive understanding of how the CtsR repressor is regulated in response to heat shock, this study aimed to determine the cryo-EM structures of *B. subtilis* McsAZ2-McsB and its complex with CtsR. Additionally, we examined CtsR binding to the operator in the presence of McsAB using anisotropy assays with fluorescently labeled operator DNA. Our structural and biochemical analyses demonstrate that McsAB antagonizes CtsR binding to the operator under heat shock conditions through autophosphorylation.

## RESULTS

### Structure determination of B. subtilis McsAZ2-McsB and McsAZ2-McsB-CtsR complexes

To investigate the role of McsAB during the heat shock response, we purified *B. subtilis* McsAZ2- McsB and McsAZ2-McsB-CtsR complexes and determined their structures using cryo-EM. The McsAZ2-McsB complex was coexpressed in *Escherichia coli* and purified through immobilized metal affinity chromatography (IMAC), followed by size-exclusion chromatography (SEC) (**Supplementary Fig. 1**). We collected 5,903 cryo-EM images of the purified McsAZ2-McsB using a 300 kV cryo-transmission electron microscope (**Supplementary Fig. 1**). Through two-dimensional (2D) and three-dimensional (3D) classification of the particles, two classes of cryo-EM density maps were reconstructed. The class 1 density map, representing a rod-shaped conformation, was reconstructed from 228,764 particles at an overall resolution of 2.89 Å. The class 2 density map, featuring two intersecting rods, was reconstructed from 266,396 particles at an overall resolution of 3.02 Å (**Supplementary Figs. 1 and 2**). Atomic models of the McsAZ2- McsB complex were built by tracing the cryo-EM density maps and refined through iterative model correction and structure refinement. The final models exhibited no outliers in the Ramachandran plot, with correlation coefficients (CCmask) of 0.81 for class 1 and 0.77 for class 2 (**Supplementary Table 1**).

The McsAZ2-McsB-CtsR complex was purified through SEC from a mixture of the purified McsAZ2- McsB complex and excess CtsR. Two sets of cryo-EM images, comprising 12,050 and 2,498 images, were collected from independently purified complexes. The cryo-EM density map for dataset 1 was reconstructed to an overall resolution of 3.56 Å from 15,254 particles (**Supplementary Figs. 3 and 5**), and the density map for dataset 2 was reconstructed to an overall resolution of 3.38 Å from 36,545 particles (**Supplementary Figs. 4 and 5**). The final models of the McsAZ2-McsB-CtsR complexes were built without any Ramachandran plot outliers, with correlation coefficients (CCmask) of 0.76 for dataset 1 and 0.82 for dataset 2 (**Supplementary Table 1**).

### Overall structure of the McsAZ2-McsB heterotetramer

The Class 1 cryo-EM structure of McsAZ2-McsB reveals a rod-shaped heterotetramer comprising a McsB dimer and two McsAZ2 monomers (**Fig. 1A and 1B**). In this structure, McsB contains a kinase domain (McsBKD; residues 2–261) and a dimerization domain (McsBDD; residues 262–354) (**Fig. 1C and Supplementary Fig. 6**). McsBKD is characterized by a central β-sheet composed of nine β-strands arranged antiparallel in the topological order S2, S3, S4, S5, S6, S1, S9, S11, and S10. These strands are interspersed with nine helices (H1–H9) and a small β-sheet (S7 and S8 strands) (**Supplementary Fig. 6**). Meanwhile, McsBDD forms a helical bundle consisting of five helices (H10–H14) and interacts with McsBKD to form a rigid globular structure (**Fig. S6**). McsAZ2 contains four helices (α1–α4) and coordinates a zinc atom through four cysteine residues (C81, C84, C99, and C102) (**Fig. 1C and Supplementary Fig. 7A and 7B)**.

**Figure 1.**
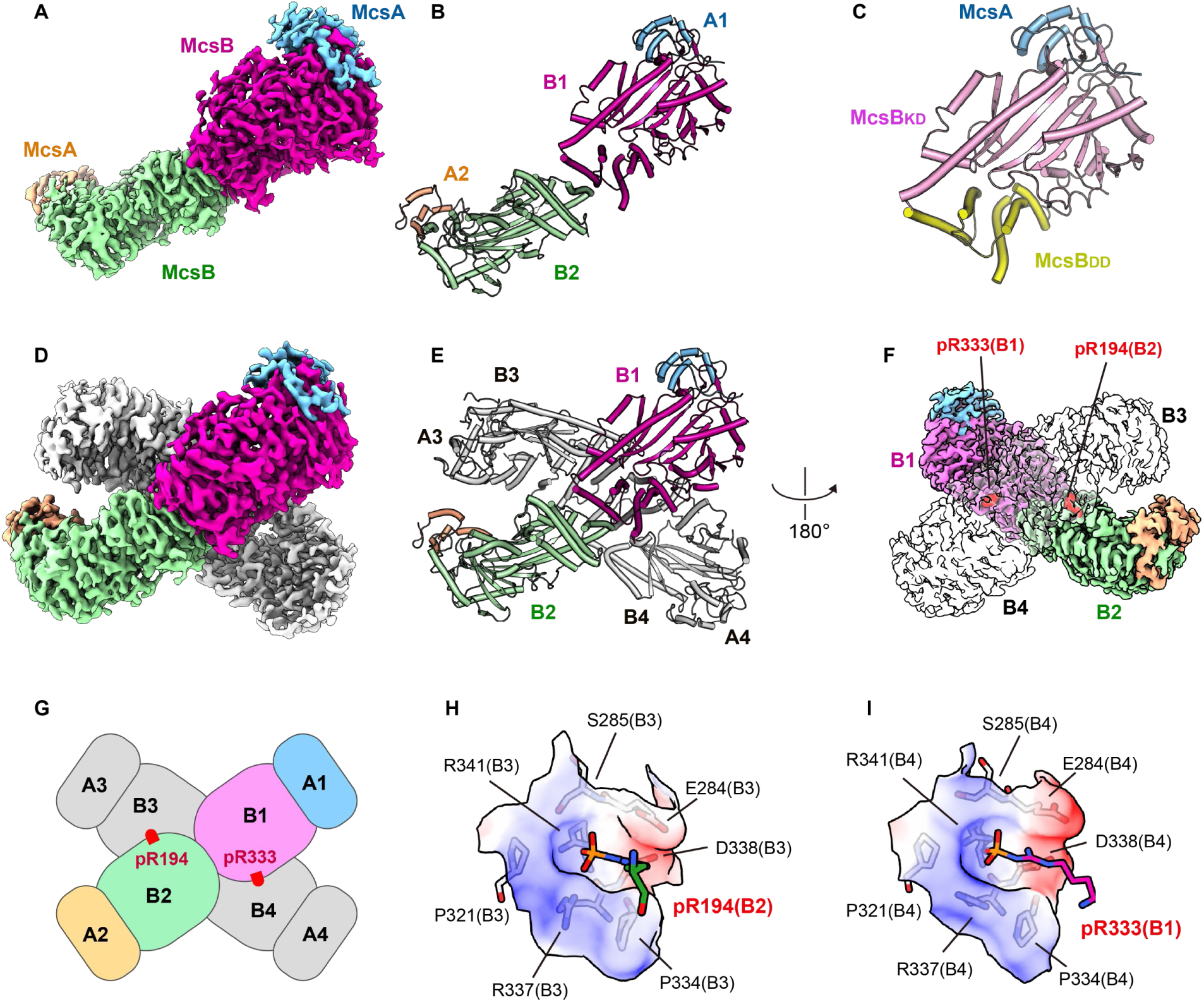
Cryo-EM structure of the McsAZ2-McsB complex. (A–C) Class 1 cryo-EM structure of the McsAZ2-McsB complex. (A, B) Cryo-EM density map of the McsAZ2-McsB heterotetramer (A) and the corresponding structural model (B). Each chain is represented in a different color. (C) Ribbon diagram depicting the structure of a McsAZ2-McsB heterodimer unit. (D–I) Class 2 cryo- EM structure of the McsAZ2-McsB complex, showing two heterotetramers intersecting asymmetrically. (D, E) Cryo-EM density map of the McsAZ2-McsB heterooctamer (D) and the corresponding structural model (E). Each chain of one heterotetramer is represented in a different color, and the intersecting tetramer is displayed in gray. McsB chains are labeled B1–B4. (F) Cryo-EM density map showing the positions of pArg residues, with pR194 in chain B2 and pR333 in chain B1 highlighted in red. The heterotetramer interacting with the pArg residues is shown transparently. (G) Cartoon representation of the positions of pR194 and pR333. Chains A1–A4 represent McsA, and chains B1–B4 represent McsB. (H, I) Interactions between pArg residues and positively charged pockets of McsB chains. Pockets are illustrated with partially transparent solvent-accessible surfaces along with stick models of the relevant residues. Electrostatic potential is shown on a color scale ranging from red (−10 kcal/mol·e) to blue (+10 kcal/mol·e). (H) Interactions between pR194 and the positively charged pocket of McsB chain B3. (I) Interactions between pR333 and the positively charged pocket of McsB chain B4.

**Figure 2.**
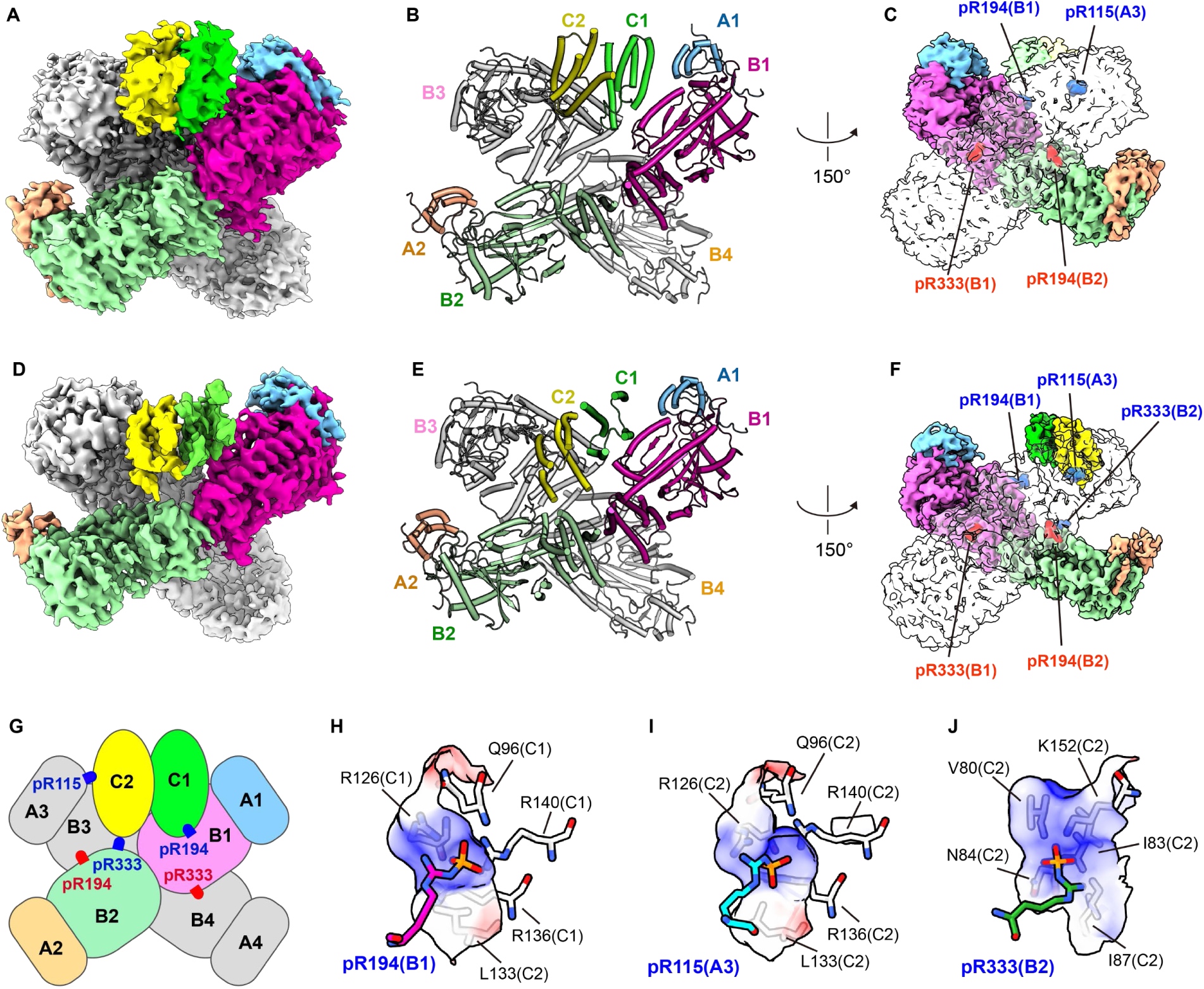
Cryo-EM structure of the McsAZ2-McsB-CtsR complex. (A–C) Cryo-EM structure of the McsAZ2-McsB-CtsR complex reconstructed from Dataset 1. (A, B) Cryo-EM density map (A) and corresponding structural model shown as a ribbon diagram (B). The chains of one McsAZ2- McsB heterotetramer and the CtsR dimer are represented in different colors, and the other McsAZ2-McsB heterotetramer is shown in gray. Chains A1–A4, B1–B4, and C1–C2 represent McsA, McsB, and CtsR, respectively. (C) Cryo-EM density map highlighting the positions of pArg residues in the Dataset 1 cryo-EM structure. pArg residues involved in the assembly of the McsAZ2-McsB oligomer are labeled in red, and the residues interacting with the CtsR dimer are shown in blue. (D–J) Cryo-EM structure of the McsAZ2-McsB-CtsR complex reconstructed from Dataset 2. Panels (D–F) are presented in the same manner as panels (A–C). (D, E) Cryo-EM density map (D) and corresponding structural model (E). (F) Cryo-EM density map highlighting the positions of pArg residues. (G) Cartoon representation showing the positions of pArg residues in the Dataset 2 structure. (H–J) Interactions between pArg residues of McsAB and surface pockets of CtsR. Pockets are depicted using partially transparent solvent-accessible surfaces along with stick models of the relevant residues. Electrostatic potential is represented by a color scale ranging from red (−10 kcal/mol·*e*) to blue (+10 kcal/mol·*e*). (H) Interactions between pR194 from McsB chain B1 and the CtsR pocket. (I) Interactions between pR115 from McsA chain A3 and the CtsR pocket. (J) Interaction between pR333 from McsB chain B2 and the CtsR pocket.

In the Class 1 cryo-EM structure, the two McsB monomers bind symmetrically through their dimerization domains, with McsAZ2 positioned on both sides of the McsB dimer (**Fig. 1A–1C**). The two McsAZ2-McsB heterodimers observed in the structure are nearly identical, with a root-mean-square deviation (RMSD) of 0.9 Å for 397 Cα atoms (**Supplementary Fig. 7C**). This cryo-EM structure of McsAZ2-McsB heterotetramer shares an overall conformation with its crystal structures^13,15^. Superimposition with the crystal structure of *B. subtilis* McsB (PDB ID: 6TV6)^13^ yielded an RMSD value of 1.5 Å for the 691 Cα atoms (**Supplementary Fig. 7D**). Additionally, superimposition of this structure on the crystal structure of *B. subtilis* McsA-McsBKD (PDB ID: 8WTC)^15^ yielded an RMSD value of 0.9 Å for 279 Cα atoms (**Supplementary Fig. 7E**).

### pArg-mediated assembly of the McsAZ2-McsB oligomer

The Class 2 cryo-EM structure of the McsAZ2-McsB complex consists of two rod-shaped tetramers (**Fig. 1D and 1E**). One of these tetramers closely resembles the Class 1 cryo-EM structure, with an RMSD of 1.2 Å for 800 Cα atoms (**Supplementary Fig. 8A–8C**). In contrast, the other tetramer exhibits a slightly more concave shape, resulting in a higher RMSD value (2.5 Å for 796 Cα atoms) (**Supplementary Fig. 8A–8C**). Together, these two tetramers form an X-shaped structure that intersects asymmetrically (**Fig. 1D and 1E**).

The interactions between the two McsAZ2-McsB tetramers are mediated only by two pArg residues in McsB (**Fig. 1F and 1G**). The pArg residues, specifically pR194 and pR333, bind to the same binding pockets in different McsB subunits. These binding pockets are positively charged^22^, and the phosphate groups of both pArg residues play a key role in this interaction (**Fig. 1H and 1I**). Specifically, the phosphate group of pR194 in McsB (chain B2) interacts with the nitrogen atoms of R337, R341, and S285 in McsB (chain B3) (**Fig. 1H**). Similarly, the phosphate group of pR333 in McsB (chain B1) interacts with the nitrogen atoms of residues R341 and S285 in McsB chain B4 (**Fig. 1I**). No other interactions between the two tetramers were observed, indicating that the autophosphorylation of McsB at residues R194 and R333 is crucial for the assembly of McsAZ2- McsB oligomers. This pArg-mediated interaction has also been observed in the crystal structure of *B. subtilis* McsB octamer (PDB ID: 6TV6)^13^. In that structure, pR194 binds to the pocket of a neighboring McsB subunit, contributing to the formation of a tetrameric ring structure. The binding pocket involved is identical to that involved in the pArg-mediated interaction in the class 2 cryo-EM structure of McsAZ2-McsB.

In the cryo-EM structure of *Staphylococcus aureus* McsAB (PDB ID: 8GQD)^16^, two rod-shaped McsAB tetramers form a symmetric dimer (**Supplementary Fig. 8D and 8E**). This structure differs from the asymmetrically intersecting structure of the *B. subtilis* McsAZ2-McsB complex (**Supplementary Fig. 8F and 8G**). This difference is likely attributable to variations in the Arg residues mediating the interactions between the two tetramers. *S. aureus* McsB contains R184, which corresponds to R194 in *B. subtilis* McsB; however, it lacks an Arg residue that corresponds to R333 in *B. subtilis* McsB (**Supplementary Fig. 6B**). Instead, this residue is substituted by D318 in *S. aureus*. The pArg-mediated interaction observed in *B. subtilis* McsAZ2-McsB complex is absent in *S. aureus* McsAB.

### pArg-mediated CtsR binding of the McsAZ2-McsB complex

We determined two types of cryo-EM structures for the McsAZ2-McsB-CtsR complex (**Supplementary Figs. 3–5**). Each structure consists of two McsAZ2-McsB heterotetramers and one CtsR dimer, resulting in a stoichiometric ratio of 4:4:2 for McsAZ2, McsB, and CtsR in the complex (**Fig. 2**). The overall shape of the McsAZ2-McsB complex in these cryo-EM structures resembles the Class 2 cryo-EM structure of the McsAZ2-McsB complex. The structures from dataset 1 and dataset 2 of the McsAZ2-McsB-CtsR complex were superimposed onto the Class 2 structure of the McsAZ2-McsB complex, yielding RMSD values of 2.6 Å for 1,395 Cα atoms and 2.0 Å for 1,565 Cα atoms, respectively (**Supplementary Fig. 9A and 9B**). Interactions between the two McsAZ2-McsB heterotetramers are mediated by pR194 and pR333, as in the Class 2 McsAZ2-McsB structure (**Figs. 1F, 1G, 2C, 2F, and 2G, and Supplementary Fig. 9C–9F**). This structural comparison reveals that the asymmetric, X-shaped structure of the McsAZ2-McsB complex remains largely unchanged upon CtsR binding.

In the cryo-EM structure of the McsAZ2-McsB-CtsR complex, CtsR binds to the groove formed between the intersecting McsAZ2-McsB tetramers (**Fig. 2**). Although full-length CtsR was purified with McsAZ2-McsB for structural analysis, the N-terminal DNA-binding domain (NTD) was not observed in the cryo-EM density map; only the dimer of the C-terminal domain (CTD) was observed (**Fig. 2A and 2D**). Consistent with this, CtsRCTD coeluted with McsAZ2-McsB during SEC, whereas CtsRNTD did not (**Supplementary Fig. 10A–10D**), indicating that CtsRCTD directly interacts with the McsAZ2-McsB complex. In the structures of the McsAZ2-McsB-CtsR complex, CtsRCTD consists of five helices (h1–h5) and forms a dimer (**Supplementary Fig. 10E**). No significant structural differences were observed between the CtsRCTD dimers from the two cryo-EM structures, with an RMSD of 1.5 Å for 148 Cα atoms (**Supplementary Fig. 10F**).

The interactions between the McsAZ2-McsB complex and the CtsR dimer are mediated by pArg residues (**Fig. 2G–2J and Supplementary Fig. 10G–10I**). In the Dataset 1 structure, the McsAZ2- McsB complex contacts both sides of the CtsR dimer through two pArg residues positioned on either side of the binding groove. Specifically, pR194 of McsB (chain B1) interacts with R136 and D137 of CtsR (chain C1) (**Supplementary Fig. 10H**). Meanwhile, pR115 of McsA (chain A3) interacts with R126 and Q96 of CtsR (chain C2) (**Supplementary Fig. 10I**). These observations indicate that both pArg residues are involved in interactions with the same two binding pockets of the CtsR dimer. In the Dataset 2 structure, pR333 of McsB participates in an additional interaction with CtsR, along with pR194 of McsB and pR115 of McsA. Specifically, pR194 of McsB (chain B1) binds to R126, R136, R140, and Q96 of CtsR (chain C1**) (Fig. 2H)**. pR115 of McsA (chain A3) interacts with R126 and R136 of CtsR (chain C2) (**Fig. 2I**). Additionally, pR333 of McsB (chain B2) interacts with N84 and K152 of CtsR (chain C2) **(Fig. 2J)**. The orientation of CtsR in the superimposed structures from Datasets 1 and 2 differs by approximately 75 degrees due to the additional interaction between pR333 of McsB and CtsR in Dataset 2. The interaction with pR333 appears to stabilize CtsR binding to McsAB by providing an additional interaction site. Overall, these results suggest that R194 of McsB and R115 of McsA are essential for CtsR binding, and R333 of McsB also contributes to enhancing overall binding stability.

### McsAB hijacks CtsR from the operator in response to heat shock

To investigate the effect of McsAB on the binding of the CtsR repressor to its operator, we performed fluorescence anisotropy experiments using 6-FAM-labeled DNA containing the CtsR operator. The operator showed stable anisotropy values for 6 h at both 30 °C and 50 °C, indicating that the DNA remained unchanged during this period (**Fig. 3A and 3B**). In contrast, anisotropy values increased when CtsR was mixed with the operator, demonstrating that CtsR binds to the operator. At 30 °C, the anisotropy in the presence of CtsR remained constant throughout the 6-h period (**Fig. 3A**). However, at 50 °C, the anisotropy gradually decreased during the same period (**Fig. 3B**). This indicates that CtsR maintains its binding to DNA under normal temperature conditions but slowly dissociates from the operator under heat shock conditions. Therefore, it is unlikely that CtsR functions as a heat shock sensor that requires a rapid stress response (**Fig. 3B**).

**Figure 3.**
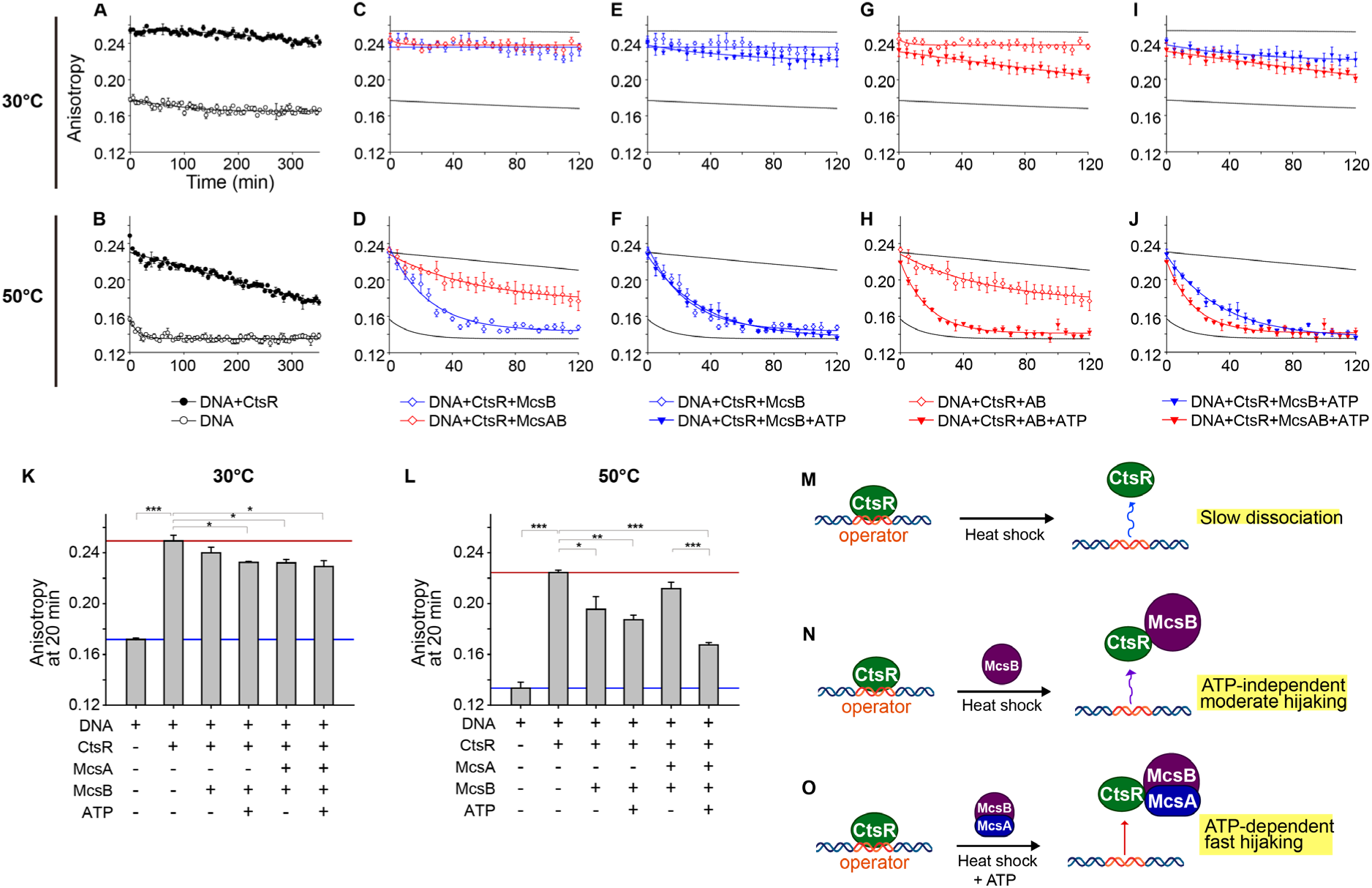
McsAB hijacks CtsR from its operator in response to heat shock. (A, B) Anisotropy measurements of 6-FAM-labeled DNA with the CtsR operator sequence. Measurements were conducted for 6 h in the presence or absence of CtsR at 30 °C (A) and 50 °C (B). (C, D) Anisotropy measurements of the CtsR-operator complex in the presence of McsB or McsAB. Anisotropy was measured for 2 h at 30 °C (C) and 50 °C (D). (E, F) Anisotropy of a mixture consisting of CtsR, its operator, and McsB in the presence or absence of ATP. The anisotropy was measured for 2 h at 30 °C (E) and 50 °C (F). (G, H) Anisotropy of a mixture consisting of CtsR, its operator, and McsAB in the presence or absence of ATP, measured for 2 h at 30 °C (G) and 50 °C (H). (I, J) Anisotropy of a mixture of CtsR, its operator, and ATP in the presence of McsB or McsAB, measured for 2 h at 30 °C (I) and 50 °C (J). (K, L) Bar graphs showing anisotropy measurements at 20 min at 30 °C (K) and 50 °C (L). Data are presented as the mean ± standard error from three independent experiments, each with three replicates. Asterisks indicate statistical significance: \**p* < 0.05, \*\**p* < 0.01, and \*\*\**p* < 0.001. (M–O) Cartoon models depicting the anisotropy results. (M) CtsR gradually dissociates from its operator over 6 h at 50 °C. (N) McsB hijacks CtsR from its operator in an ATP-independent manner during heat shock. (O) McsAB hijacks CtsR from its operator more rapidly than McsB in an ATP-dependent manner during heat shock.

Next, we examined changes in the anisotropy of the CtsR-operator complex in the presence of either McsAB or McsB (**Fig. 3C, 3D, 3K, and 3L**). To prepare the McsAB complex, individual McsA and McsB proteins were purified and mixed in a 1:1 molar ratio. At 30 °C, anisotropy values remained relatively stable, indicating that neither McsB nor McsAB affects the stability of the CtsR- operator complex or interacts with it under normal conditions (**Fig. 3C and 3K**). In contrast, at 50 °C, anisotropy values decreased significantly, with a more rapid decline observed in the presence of McsB compared to McsAB (**Fig. 3D and 3L**). These results suggest that McsB disrupts the CtsR-operator complex under heat shock conditions in an ATP-independent manner.

We also investigated the effect of ATP on the activity of McsB and McsAB in hijacking CtsR from its operator (**Fig. 3E–3L**). The addition of ATP did not alter the anisotropy of the CtsR-operator complex by McsB at either 30 °C or 50 °C, indicating that the CtsR-hijacking activity of McsB is independent of ATP (**Fig. 3E and 3F**). In contrast, the CtsR-hijacking activity of McsAB was enhanced in the presence of ATP (**Fig. 3G and 3H**). At 30 °C, the presence of ATP caused a slight reduction in the anisotropy of the CtsR-operator complex by McsAB compared to when ATP was absent (**Fig. 3G and 3K**). At 50 °C, this reduction was more rapid and greater than that observed with McsB (**Fig. 3H and 3L**). These results show that McsAB hijacks CtsR from the operator more efficiently than McsB in the presence of ATP. Considering that McsA and McsB are encoded in the same operon and that ATP is available in the bacterial cytoplasm, it is likely that McsAB, rather than McsB, is responsible for removing CtsR from the operator in response to heat shock.

We further investigated the CtsR-hijacking activity of the McsAZ2-McsB complex (**Supplementary Fig. 11**). In this experiment, McsAZ2 and McsB were coexpressed and purified as a complex. The McsAZ2-McsB complex did not alter the anisotropy of the CtsR-operator complex at 30 °C; however, it significantly reduced the anisotropy at 50 °C (**Supplementary Fig. 11A–11J**). This indicates that the McsAZ2-McsB complex disrupts the CtsR-operator complex, similar to McsAB. However, the CtsR-hijacking activity of McsAZ2-McsB was not affected by ATP, in contrast to McsAB (**Supplementary Fig. 11J**). This difference appears to be due to the McsAZ2-McsB complex being autophosphorylated during the coexpression in *E. coli*.

### pArg residues in McsA and McsB are crucial for CtsR hijacking from the operator during heat shock

Since McsAB functions as a kinase, the enhanced hijacking of CtsR by McsAB in the presence of ATP is likely related to phosphorylation. Notably, cryo-EM structures of the McsAZ2-McsB- CtsR complex show that pArg residues of McsAB mediate the binding of CtsR. To investigate whether McsAB hijacks CtsR from the operator through autophosphorylation, we performed anisotropy experiments using McsAB mutants. In this experiment, R115 of McsA and R194 of McsB, which mediate CtsR binding, were mutated to alanine. Native McsA, native McsB, McsA(R115A), and McsB(R194A) were purified individually. Combinations of McsAB mutants were prepared by mixing purified McsA and McsB proteins in a 1:1 molar ratio prior to the reaction. Compared to native McsAB, three McsAB mutant proteins (McsA(R115A)-McsB, McsA-McsB(R194A), and McsA(R115A)-McsB(R194A)) showed a slight increase in anisotropy at both 30°C and 50°C in the absence of ATP, as well as at 30°C in the presence of ATP; however, these changes were not statistically significant (**Fig. 4A–4C**). In contrast, at 50 °C in the presence of ATP, the McsAB mutants resulted in a significant increase in anisotropy (**Fig. 4D**). This suggests that the arginine residues involved in CtsR binding contribute to the CtsR-hijacking activity of McsAB.

**Figure 4.**
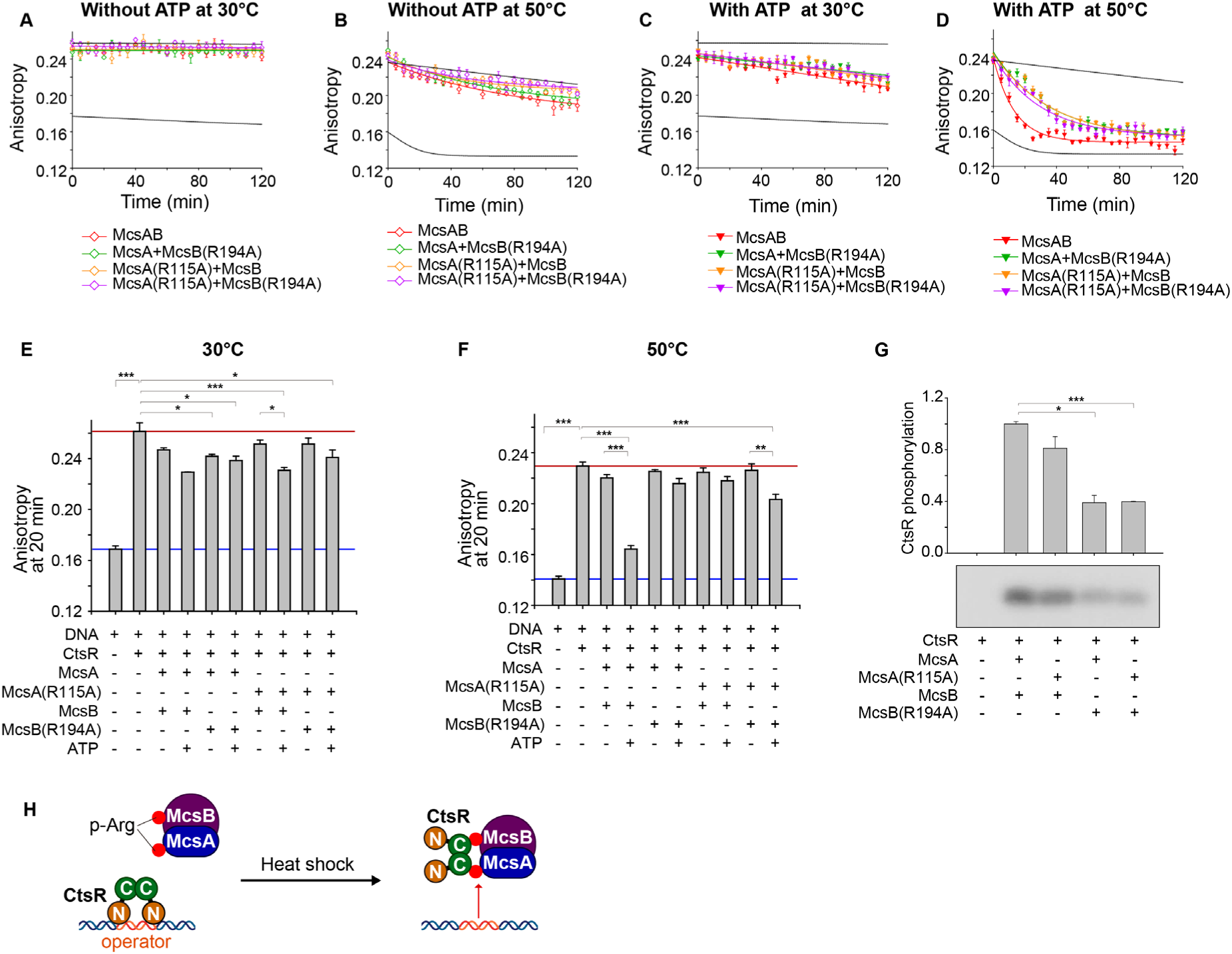
Autophosphorylation-dependent hijacking of CtsR by McsAB during heat shock. (A, B) Anisotropy measurements of the CtsR-operator complex in the presence of McsAB or its mutants. Anisotropy was measured without ATP for 2 h at 30 °C (A) and 50 °C (B). (C, D) Anisotropy measurements of the CtsR-operator complex in the presence of McsAB or its mutants. Anisotropy was measured with ATP for 2 h at 30 °C (C) and 50 °C (D). (E, F) Bar graphs showing anisotropy values at 20 min at 30 °C (E) and 50 °C (F). Data are presented as the mean ± standard error from three independent experiments, each with three replicates. (G) Kinase activity of McsAB and its mutants, assessed by measuring CtsR phosphorylation. Data are presented as mean ± standard error from three independent experiments. Asterisks in panels (E–G) denote *p* values: \**p* < 0.05, \*\**p* < 0.01, and \*\*\**p* < 0.001. (H) Cartoon model depicting autophosphorylation- dependent CtsR-hijacking of McsAB.

The McsAB mutants exhibited partial CtsR-hijacking activity at 50 °C in the presence of ATP, indicating that mutations in the arginine residues do not completely abolish the activity (**Fig. 4D**). Similarly, mutations in the McsAZ2-McsB complex reduced its CtsR-hijacking activity but did not abolish it (**Supplementary Fig. 11**). These results suggest that other factors, such as CtsR phosphorylation or its interaction with the active site of McsAB, may contribute to the effective hijacking of CtsR from the operator, in addition to the autophosphorylation of McsAB. Supporting this, the McsAZ2(R115A)-McsB(R194A) mutant coeluted with either CtsR or CtsRCTD during SEC. The elution volumes of CtsR and CtsRCTD matched those of McsB(R194A) but not McsA(R115A), suggesting that CtsR interacts with McsB when the pArg-mediated binding site is disrupted **(Supplementary Fig. 12A–12F)**. To further investigate the interaction between the McsA(R115A)-McsB(R194A) mutant and CtsR, we reconstructed a cryo-EM density map of the McsA(R115A)-McsB(R194A)-CtsR complex. The overall shape of the cryo-EM density map closely resembled the rod-shaped McsAZ2-McsB tetramer, with no observation of an intersecting X-shaped structure. This is likely due to the R194A mutation in McsB, which affects the assembly of the McsAZ2-McsB oligomer (**Supplementary Fig. 12G and 12H**). Additionally, no density corresponding to CtsR was observed in this cryo-EM structure, indicating that CtsR may interact with McsB through a nonspecific binding site in the absence of a pArg-mediated binding site. These results suggest that CtsR can bind to McsB of the McsAB complex in a pArg-independent and relatively nonspecific manner, in addition to the pArg-dependent binding site of McsAB.

Next, we assessed the kinase activity of the McsAB mutants regarding CtsR phosphorylation (**Fig. 4G**). Compared to native McsAB, the McsA(R115A)-McsB mutant showed a slight decrease in activity, approximately 20%. In contrast, the McsA-McsB(R194A) mutant exhibited a significant reduction in activity, exceeding 60%. The activity levels of the McsA(R115A)-McsB(R194A) mutant were similar to that of the McsA-McsB(R194A) mutant. The changes in activity caused by the McsAB mutations closely correlated with the alterations in CtsR-hijacking activity. This suggests that R115 in McsA and R194 in McsB, which are located away from the kinase active site, are required for effective CtsR phosphorylation, as well as CtsR hijacking during heat shock.

## DISCUSSION

Covalent modifications of proteins regulate their activity and ligand binding by altering the steric and polar properties of amino acids^23^. Phosphorylation is the most common of these modifications. Amino acids with reactive side chains, such as hydroxyl, amino, carboxyl, and sulfhydryl groups, are targeted for phosphorylation^24^. Notably, the phosphorylation of arginine results in a significant change in the surface properties of proteins, transforming a positive charge into a negative one^25^. In gram-positive bacteria, pArg has been shown to mediate the self-assembly of McsB into a chamber-shaped octamer^22^ and to promote substrate binding to ClpCP for degradation^14^. In our study, cryo-EM structures of the *B. subtilis* McsAZ2-McsB and its complex with CtsR reveal that pArg residues are essential for the assembly of the McsAB oligomer and the binding of CtsR to McsAB. Furthermore, these pArg residues influence the kinase activity of McsAB. Although the pArg-dependent CtsR-binding site is located in a different region from the enzymatically active site, mutations in the Arg residues (R115A of McsA or R194A of McsB) reduced the kinase activity of McsAB. This suggests that autophosphorylation is essential for the molecular function of McsAB.

Pull-down assays of the McsAZ2-McsB-CtsR complex have shown that McsAB can bind to CtsR at room temperature^15^. However, McsAB did not significantly alter the anisotropy of the CtsR- operator complex at 30 °C, indicating that it does not effectively displace CtsR from the operator under normal temperature conditions. McsAB disrupts the CtsR-operator complex during heat shock. The mechanism by which McsAB more effectively disrupts the CtsR-operator interaction under heat shock conditions remains unclear. This may be because of the effects of elevated temperature and potential conformational changes in CtsR. CtsR dissociates slowly from the operator at 50 °C, suggesting that its interactions with the operator are weakened under heat shock conditions^26^. In this state, the CtsRCTD, responsible for CtsR dimerization, may undergo slight conformational changes that promote operator dissociation and McsAB binding. Consequently, the weakening of the interaction between CtsR and the operator by heat shock is likely further facilitated by McsAB, allowing a rapid response to heat shock.

Our anisotropy experiments demonstrate that the hijacking of CtsR from its operator by McsAB depends on pArg residues and heat shock conditions. This indicates that autophosphorylated McsAB can stimulate heat shock transcription by disrupting the interaction between CtsR and its operator in a temperature-dependent manner. However, mutations in the Arg residues of McsAB did not completely abolish CtsR hijacking from the operator, although they significantly reduced its activity. Moreover, CtsR coeluted with the McsAB mutant in a pArg-independent and nonspecific manner. This suggests that pArg-mediated CtsR hijacking is not the only mechanism by which McsAB regulates its activity in response to heat shock. It appears that this mechanism works synergistically with other regulatory processes, such as CtsR phosphorylation, to rapidly and efficiently control the interaction between CtsR and the operator.

Three Arg residues required for McsAB assembly and CtsR binding (R194 and R333 in *B. subtilis* McsB, and R115 in *B. subtilis* McsA) are not fully conserved across bacterial species (**Supplementary Figs. 6B and 7B**). McsAB proteins from most species, including *S. aureus*, lack the Arg residues corresponding to R333 of McsB and R115 of McsA. Notably, *Listeria monocytogenes* McsAB is missing all three of these Arg residues. The cryo-EM structure of *S. aureus* McsAB has shown that it assembles into a symmetric dimer of heterotetramers in a pArg- independent manner, which means that it does not form a pArg-mediated CtsR-binding site. This contrasts with *B. subtilis* McsAB, which forms an asymmetrically intersecting octamer and establishes a CtsR-binding site through pArg residues. As a result, the functional mechanisms of McsAB can vary based on the presence of Arg residues in bacterial species. Therefore, further structural analyses of McsAB complexes from various bacterial species are necessary to gain a deeper understanding of the general molecular mechanism by which pArg mediates McsAB function.

## METHODS

### Plasmid preparation

To modify the expression vector, a DNA sequence encoding a 6×His tag and a Tobacco Etch Virus protease (TEVP) cleavage site was inserted between the *NcoI* and *BamHI* restriction sites of the pETDuet-1 vector (Merck Millipore). Additionally, a DNA sequence encoding a 6×His tag, thioredoxin (Trx), and a TEVP cleavage site was inserted into the pETDuet-1 and pRSFDuet-1 vectors (Merck Millipore). These vectors are designated pET-His, pET-His-Trx, and pRSF-His-Trx, respectively.

The DNAs encoding McsA (residues 1–185; UniProt ID P37569), McsB (residues 1–363; UniProt ID P37570), and CtsR (residues 1–154; UniProt ID P37568) were amplified through polymerase chain reaction using genomic DNA from *B. subtilis* strain 168. The *mcsA* gene and its fragments were inserted into the pET-22b and pRSFDuet-1 vectors (Merck Millipore, Billerica, Massachusetts, USA) to express the protein without additional tags. The gene was also inserted into the pET-His vector to express the protein with a 6×His tag. The *mcsB* gene was inserted into the pET-His-Trx and pRSF-His-Trx vectors. The *ctsR* gene and its fragments were inserted into the pET-His-Trx vector. Point mutations were introduced directly into the plasmids through site-directed mutagenesis.

### Protein expression and purification

For the expression of the individual proteins McsA, McsB, and CtsR, the plasmids pET-His-McsA, pET-His-Trx-McsB, and pET-His-Trx-CtsR were separately introduced into *E. coli* BL21-star (DE3) (Thermo Fisher Scientific, Waltham, MA, USA). For the coexpression of McsAZ2 and McsB, pRSF- McsAZ2 and pET-His-Trx-McsB were introduced together into BL21-star (DE3) cells. The plasmids containing mutant genes were introduced into BL21-star (DE3) using the same procedures as those for the plasmids carrying native genes.

The transformed cells were cultured in Luria–Bertani (LB) medium at 37 °C. Protein expression was induced by adding 0.1 mM isopropyl β-D-1-thiogalactopyranoside to the culture medium when the optical density at 600 nm reached 0.6. The cells were then cultured for 36 h at 15 °C and harvested by centrifugation at 3,000 × *g* for 15 min. The cell pellets were resuspended in buffer A (20 mM HEPES pH 7.5, 0.5 M NaCl, 0.2 mM TCEP, and 5% (v/v) glycerol) and lysed by sonication. CtsR expression was induced using 0.4 mM isopropyl β-D-1-thiogalactopyranoside, and the cells were cultured at 20 °C overnight. Cell lysates were treated with DNase I and RNase A (10 μg/mL each; Roche, Basel, Switzerland) for 30 min on ice to remove genomic DNA and RNA contamination. The mixture was clarified by centrifugation at 20,000 × *g* for 35 min at 4°C.

The proteins were purified using IMAC and SEC. For IMAC, the cell lysate was loaded onto a 5- mL HisTrap nickel-chelating column (Cytiva, Marlborough, MA, USA). The column was washed with 70 mM imidazole to remove proteins bound non-specifically to the resin. Proteins bound to the resin were eluted using a linear imidazole gradient of 0.07–1.0 M. Fractions containing the recombinant proteins were pooled and treated with TEVP at 4°C overnight. Following complete cleavage of the 6×His-Trx or 6×His tags from the target proteins, the protein solution was dialyzed in buffer A for 4 h and passed through HisPur Ni-NTA resin (Thermo Fisher Scientific) to remove the tags. The proteins were further purified through SEC using Superdex 75 and Superdex 200 preparatory grade columns (Cytiva) equilibrated with buffer A. For the purification of McsA, the protein eluted from IMAC was immediately subjected to SEC using a Superdex 200 preparatory grade column.

To prepare cryo-EM samples, McsAZ2-McsB was further purified through SEC using a Superose 6 increase 10/300 GL column (Cytiva) equilibrated with buffer B (20 mM HEPES pH 7.5, 0.5 M NaCl, 0.2 mM TCEP). For the preparation of McsAZ2-McsB-CtsR complexes, McsAZ2-McsB and CtsR were mixed at a 1:1 molar ratio, and the complex was purified using a Superose 6 Increase 10/300 GL column equilibrated with buffer B. Both McsAZ2-McsB and McsAZ2-McsB-CtsR complexes were concentrated to a final concentration of 5 mg/mL.

### Cryo-EM sample preparation and data collection

Purified McsAZ2-McsB (3 µL, 5 mg/mL) and McsAZ2-McsB-CtsR (3 µL, 5 mg/mL) complexes were applied to glow-discharged UltrAuFoil R1.2/1.3 300-mesh Au grids (SPI Supplies, West Chester, PA, USA). Grids were blotted for 3 s at a blot force of 5 using a Vitrobot Mark IV (Thermo Fisher Scientific) with humidity-saturated filter paper (Ted Pella, Redding, CA, USA) and plunge-frozen into liquid ethane. EM images were collected on a Titan Krios G4 cryo-TEM (Thermo Fisher Scientific) equipped with a Falcon 4 direct electron detector and a Selectris X energy filter (slit width: 10 eV). Micrographs were acquired at a pixel size of 0.7451 Å (nominal magnification: 165,000×), with a total electron dose of 50–60 e^−^/Å² and nominal defocus values ranging from −0.6 to −2.0 µm (**Supplementary Table 1**).

### Data processing and density map reconstruction

Image processing was performed using cryoSPARC v4.4.0^27^. Raw movies were motion-corrected using Patch Motion Correction, and contrast transfer function (CTF) parameters were estimated using Patch CTF Estimation. Micrographs were curated based on astigmatism and CTF fit resolution.

To reconstruct the cryo-EM density map of McsAZ2-McsB, a total of 5,903 micrographs were collected. Particles from these micrographs were independently picked using both the blob picker and the neural network-based picker TOPAZ^28^. Initially, 3,724,487 particles were extracted using the blob picker with a box size of 400 pixels. After multiple rounds of 2D classification, 634,035 particles were selected. Additionally, 1,435,675 particles were extracted using TOPAZ^28^. The two particle sets were merged, and duplicate and low-quality particles were removed through further rounds of 2D classification, resulting in a final dataset of 763,580 particles. These particles were categorized into five 3D classes based on an ab initio model, and non-uniform refinements^29^ were performed on the two major classes. Class 1, which displayed a rod-shaped density, was reconstructed from 228,764 particles at an overall resolution of 2.89 Å. Class 2, which exhibited a crossing shape, was reconstructed from 266,396 particles at an overall resolution of 3.02 Å.

To reconstruct the cryo-EM density map of the McsAZ2-McsB-CtsR complex, two sets of micrographs were collected. The first set consisted of 2,498 micrographs. From this set, 1,966,040 particles were extracted using the blob picker with a box size of 400 pixels, and 267,163 particles were selected after multiple rounds of 2D classification. Using TOPAZ^28^, an additional 557,676 particles were extracted as a separate dataset. The two particle sets were merged, and duplicate and low-quality particles were removed through further 2D classification, resulting in a final dataset of 448,229 particles. These particles were grouped into three 3D classes based on an ab initio model, and the major class was subjected to further refinement. The final density map was reconstructed from 15,254 particles at an overall resolution of 3.56 Å using non-uniform refinement^29^ and 3D variability analysis^30^.

The second dataset of the McsAZ2-McsB-CtsR complex consisted of 12,050 micrographs. A total of 7,576,165 particles were extracted using the blob picker with a box size of 400 pixels, and 587,085 particles were selected after multiple rounds of 2D classification. Using TOPAZ^28^, 1,872,398 particles were extracted as a separate dataset. The two particle sets were merged, and duplicate and low-quality particles were removed through additional rounds of 2D classification, resulting in a final dataset of 1,254,269 particles. These particles were categorized into four 3D classes based on an ab initio model. The major class, comprising 543,639 particles, was subjected to further refinement. The final density map was reconstructed at an overall resolution of 3.38 Å from 36,545 particles selected through non-uniform refinement^29^ and 3D variability analysis^30^.

### Atomic model building and structure analysis

The AlphaFold models^31^ of *B. subtilis* McsB and CtsR, along with the crystal structure of McsA- McsBKD (PDB ID: 8WTC)^15^, were used as templates for model building. The models were initially fitted into the cryo-EM map using ChimeraX^32^, rebuilt through manual tracing in COOT^33^, and refined using Phenix.refinement^34,35^. This process of manual fitting and refinement was repeated multiple times to obtain the final atomic model. The statistics for cryo-EM data collection and refinement are summarized in Table S1. Molecular interactions were analyzed using PISA^36^ and DIMPLOT^37^. Figures were generated using ChimeraX^32^, PyMOL^38^, and ALSCRIPT^39^.

### Fluorescence anisotropy

The CtsR operator DNA was prepared using a single-stranded DNA (5’- TGACTTTGACTATATTTGACTTTAAT-3’) with its complementary strand labeled with 5’- fluorescein phosphoramidite (5’-ATTAAAGTCAAATATAGTCAAAGTCA-(6-FAM)-3’) in buffer C (20 mM Tris-HCl pH 7.5) at a concentration of 500 nM. Protein concentrations were determined using a Nanodrop spectrophotometer (Thermo Fisher Scientific), and molar concentrations were adjusted based on the intensity of McsB in the McsAB complex, as assessed using SDS-PAGE. All proteins were prepared at a concentration of 50 μM in buffer A.

For anisotropy experiments, the reaction mixture was prepared in a 96-well microtiter plate (Corning Inc.). CtsR and operator DNA were added to 150 μL of buffer C (25 mM Tris-HCl pH 7.5, 0.5 M NaCl, and 5% (v/v) glycerol) to final concentrations of 5 nM and 1 μM, respectively. After a 10-min incubation, either 0.5 mM ATP/2.5 mM MgCl2 or an equivalent volume of buffer C was added to the mixtures. Following this, McsB, McsAB, McsAZ2-McsB, or their mutants were added to the mixture at a final concentration of 1 μM. The microtiter plate containing the reaction mixtures was then loaded into a SpectraMax iD5 microplate reader (Molecular Devices, San Jose, CA, USA) and incubated for 10 min at either 30 °C or 50 °C.

Fluorescence anisotropy was measured at excitation and emission wavelengths of 485 nm and 525 nm, respectively, with measurement intervals of 5 min. The plate was shaken for 3 s prior to each measurement. Three independent experiments, each with three replicates (nine measurements in total), were conducted. Mean values from each experiment were calculated, and the overall mean ± standard error (SE) from the three experiments was analyzed and visualized using SigmaPlot 14.0 (Systat Software Inc., San Jose, CA, USA).

### Kinase activity assay

Protein phosphorylation was assessed by measuring the amount of protein labeled with a ^32^P isotope. For the assay, 20 μL reaction mixtures were prepared, containing 5 μM McsA (or the McsA-R115A mutant), 5 μM McsB (or the McsB-R194A mutant), 20 μM CtsR, 20 mM MgCl2, 2.5 mM ATP, and 5 μCi [*γ*-^32^P]ATP (PerkinElmer, Waltham, MA, USA) in buffer A. The mixtures were incubated at 37 °C for 2 h, and the reactions were terminated by adding 2× Laemmli sample buffer. Proteins were separated using tricine-SDS-PAGE and transferred onto a polyvinylidene difluoride membrane (Merck Millipore). The membrane was exposed to autoradiography film for 24 h (Thermo Fisher Scientific). The quantity of ^32^P-labeled proteins was analyzed using ImageJ v1.5 software^40^. The mean ± standard error and *p*-values from three replicates were analyzed and visualized using SigmaPlot 14.0 (Systat Software, Inc.).

## SEC

CtsRNTD (residues 1–74) and CtsRCTD (residues 75–154) were expressed and purified using the same procedures as CtsR. Mixtures of 500 μL were prepared, containing 0.2 mM McsAZ2-McsB (or its mutant McsAZ2(R115A)-McsB(R194A)) along with an excess of CtsR (or its fragments, CtsRNTD and CtsRCTD) in buffer A. Each mixture was injected into a Superdex 200 Increase 10/300 GL column (Cytiva) equilibrated with buffer A. Proteins in the fractionated eluates were visualized using tricine-SDS-PAGE.

## ACKNOWLEDGMENTS

This research was supported by the Basic Science Research Program through the National Research Foundation of Korea (NRF), funded by the Ministry of Science and ICT (RS-2022- NR070837; RS-2024-00334946) and the Ministry of Education (RS-2023-00301974).

## AUTHOR CONTRIBUTIONS

M.A. prepared all proteins and performed anisotropy experiments; K.P. collected EM images and reconstructed cryo-EM density maps; E.K. and D.Y.K. performed kinase activity assays, built model structures, analyzed the experimental results, and wrote the manuscript with contributions from all authors

