## Supplementary information for "Structural insights into autophosphorylation-mediated heat shock response by the protein-arginine kinase McsAB"

#### Figure S1

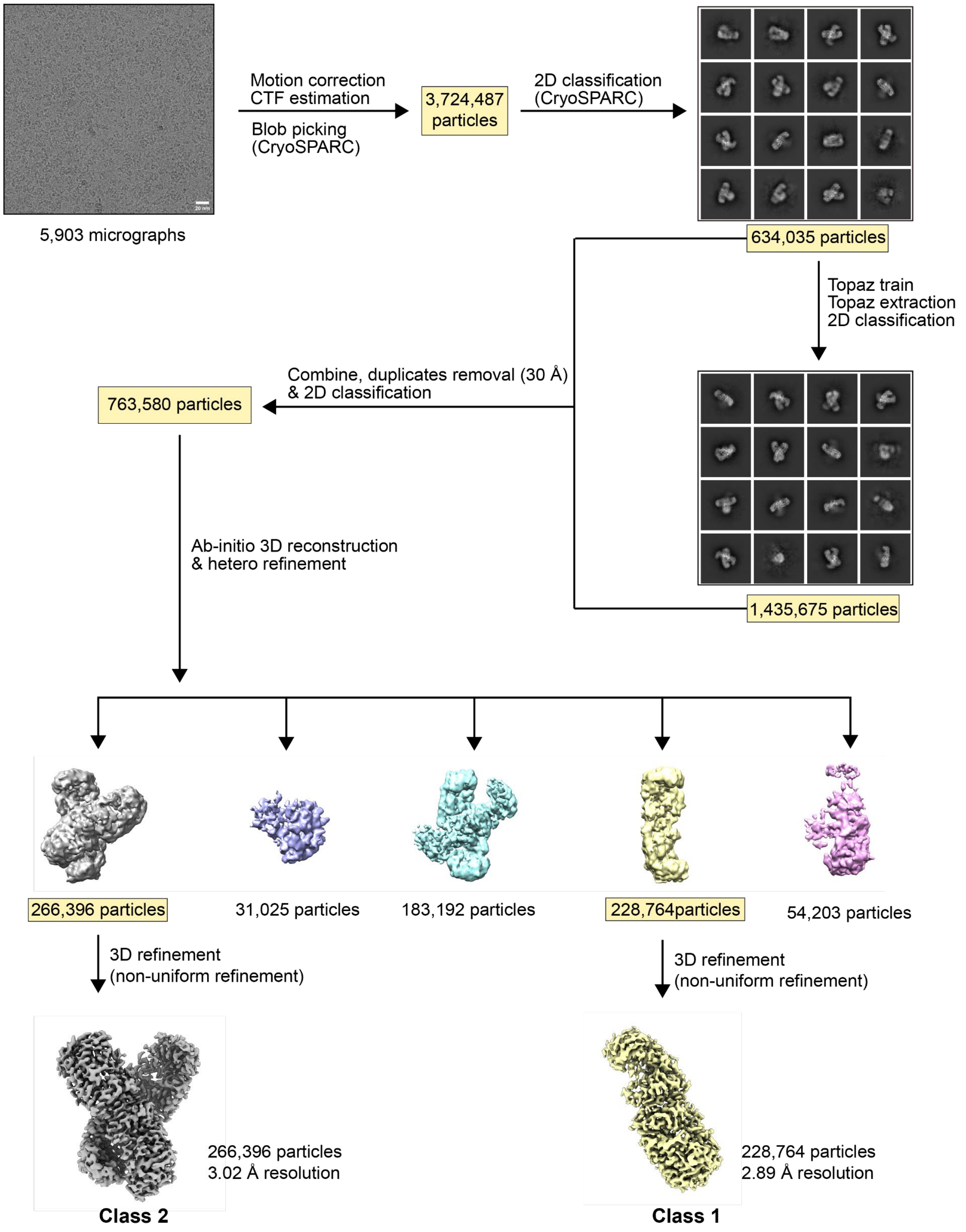


**Supplementary Figure 1. Cryo-EM data processing workflow for the McsA$_{Z2}$-McsB complex.** Scale bars on the micrograph represent 20 nm.

### Figure S2

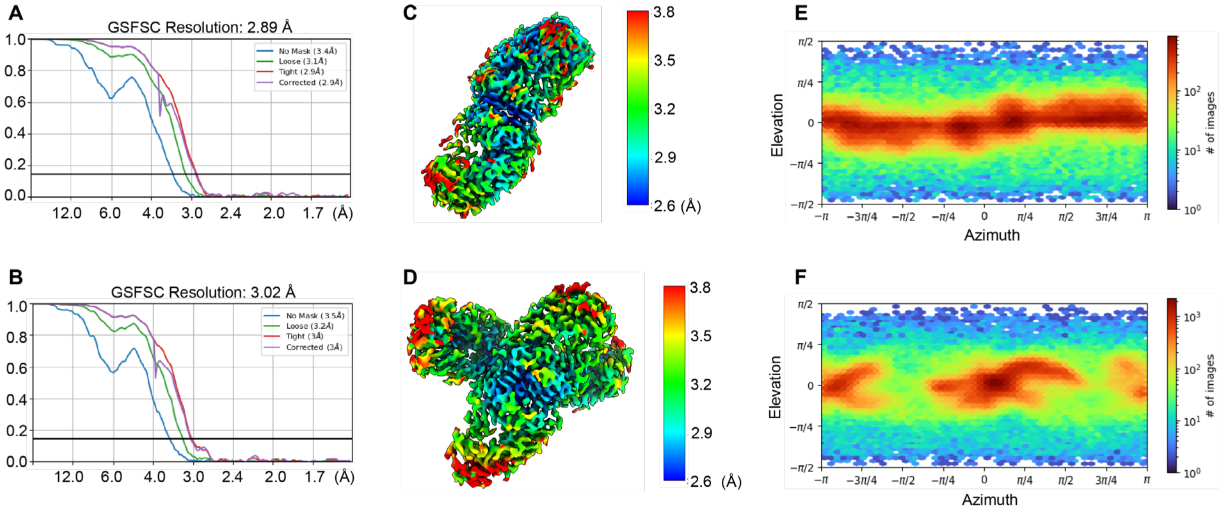


**Supplementary Figure 2. Cryo-EM data analysis of the $McsA_{Z2}$-McsB complex.** (A, B) Fourier shell correlation (FSC) curves for Class 1 (A) and Class 2 (B), with overall resolution estimated at an FSC value of 0.143. (C, D) Cryo-EM density maps of Class 1 (C) and Class 2 (D) showing the local resolution. (E, F) Angular distributions of particle projections for Class 1 (E) and Class 2 (F).

**Figure S3**

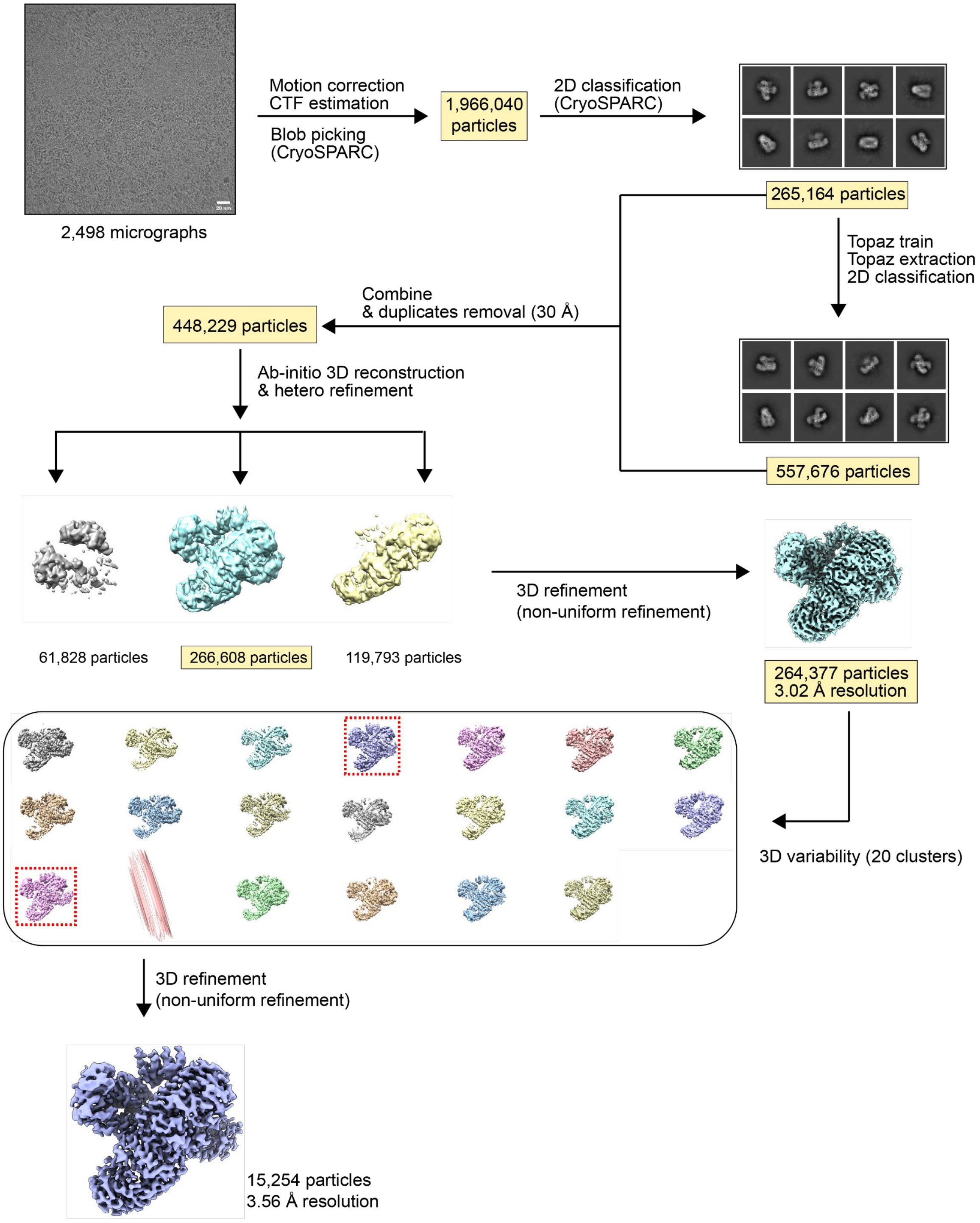


**Supplementary Figure 3. Cryo-EM data processing workflow for Dataset 1 of the McsA$_{Z2}$-McsB-CtsR complex.** Scale bars on the micrograph represent 20 nm.

### Figure S4

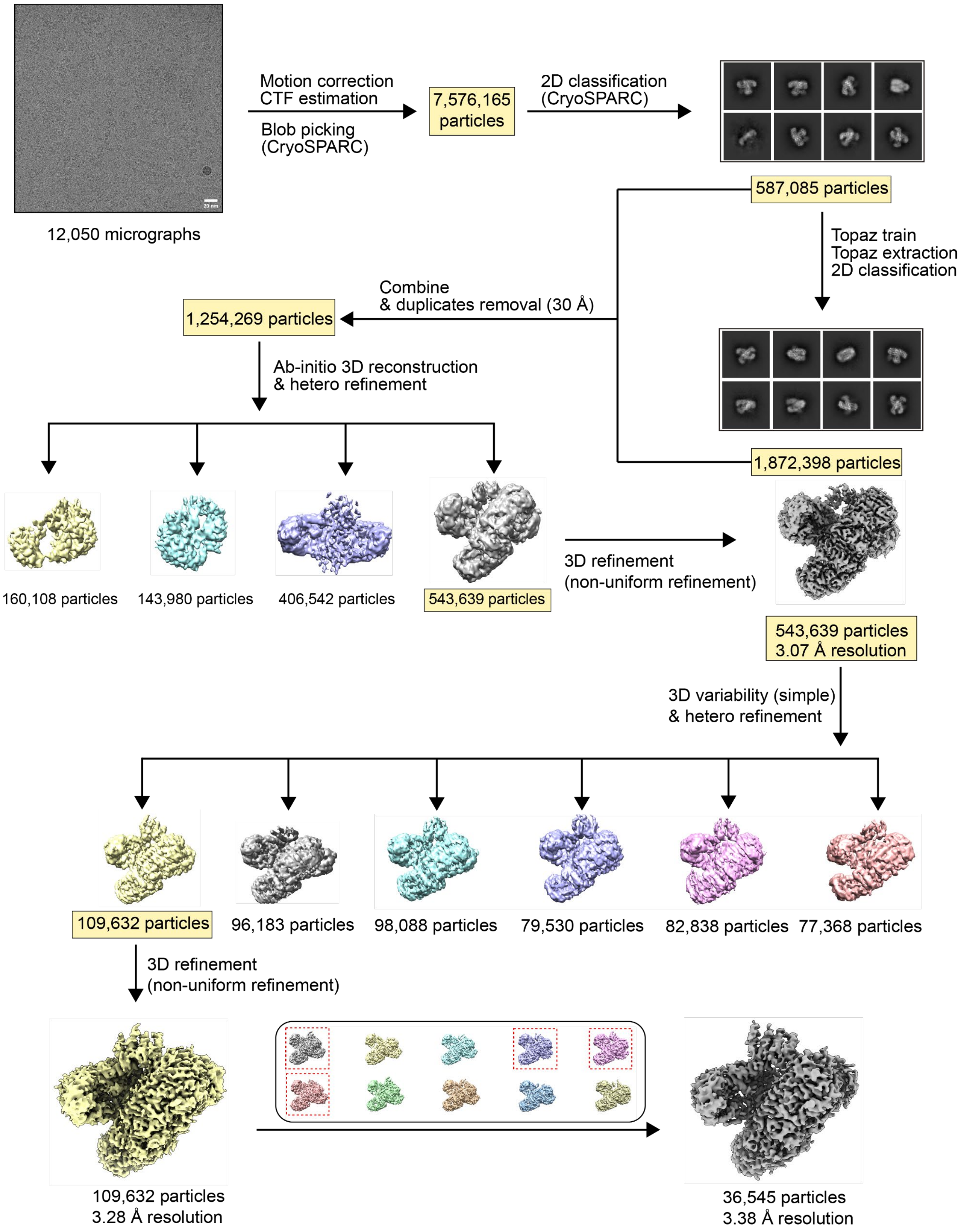


**Supplementary Figure 4. Cryo-EM data processing workflow for Dataset 2 of the McsA$_{Z2}$-McsB complex.** Scale bars on the micrograph represent 20 nm.

**Figure S5**

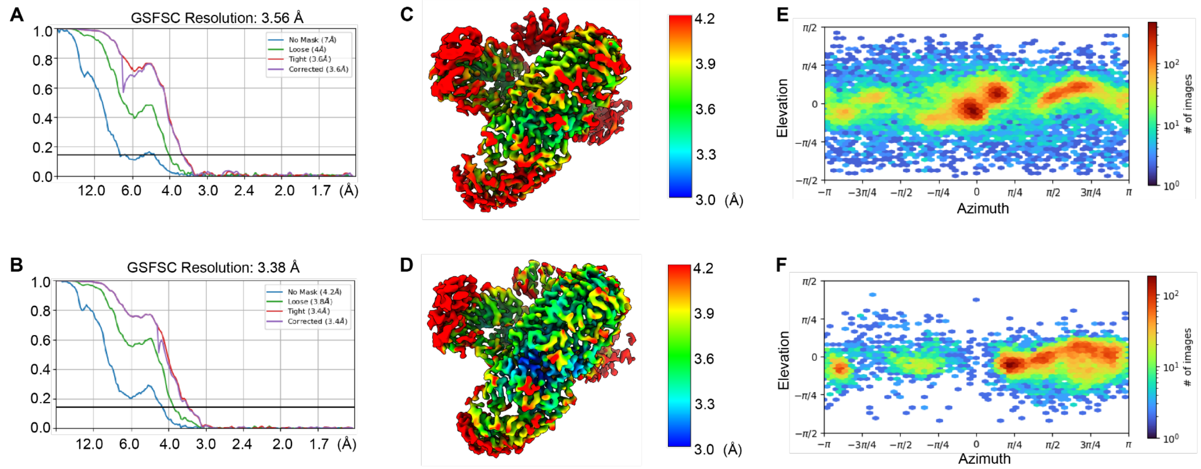


**Supplementary Figure 5. Cryo-EM data analysis of the McsA$_{Z2}$-McsB-CtsR complexes.** (A, B) Fourier shell correlation (FSC) curves for Dataset 1 (A) and Dataset 2 (B), with overall resolution estimated at an FSC value of 0.143. (C, D) Cryo-EM density maps of Dataset 1 (C) and Dataset 2 (D) showing the local resolution. (E, F) Angular distributions of particle projections for Dataset 1 (E) and Dataset 2 (F).

### Figure S6

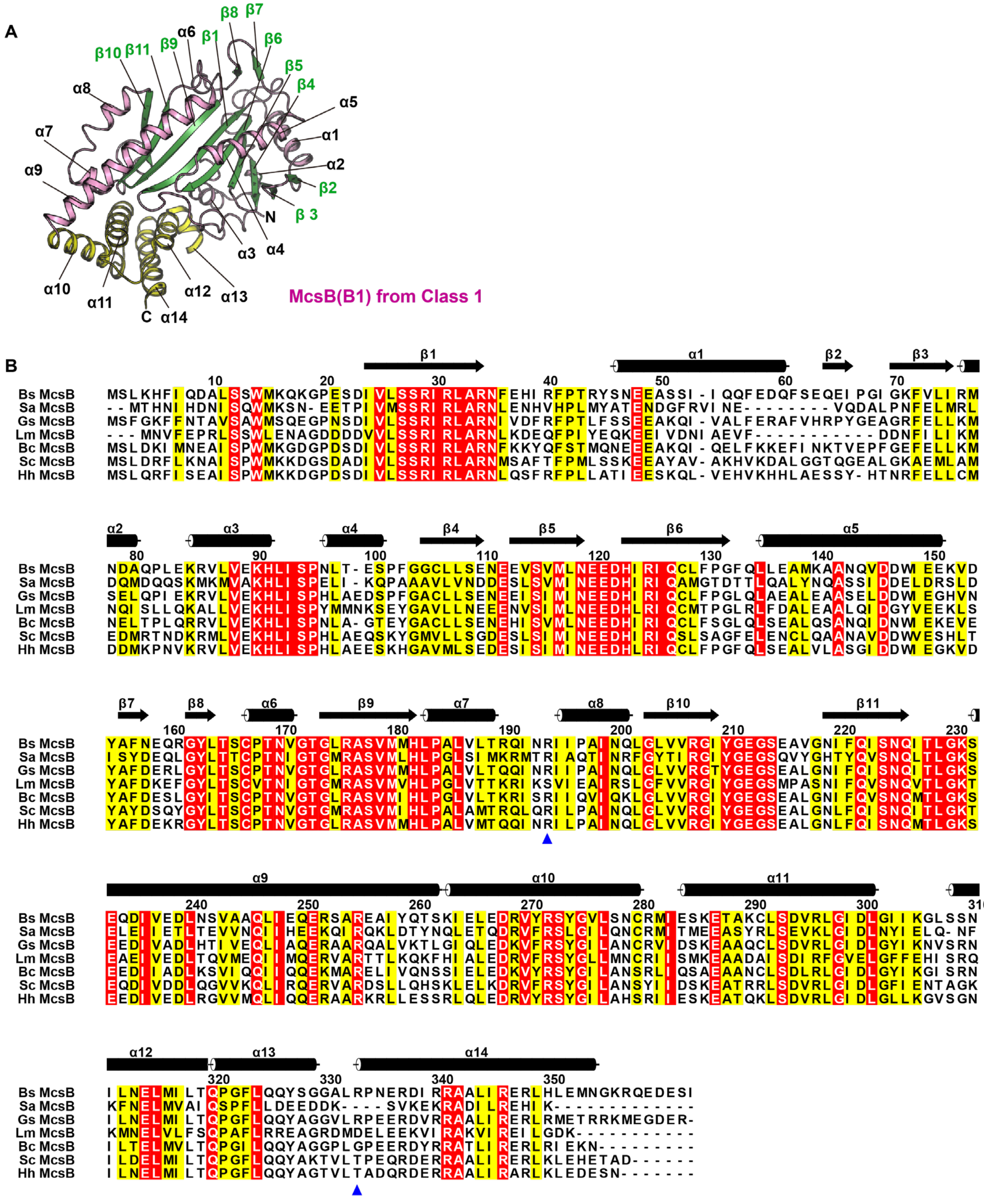


**Supplementary Figure 6. Structure of McsB in the McsA$_{Z2}$-McsB complex.** (A) Ribbon diagram of chain B1 of McsB from the Class 1 cryo-EM structure. The β-sheets and peripheral helices of the McsB$_{KD}$ and McsB$_{DD}$ are shown in green, pink, and yellow, respectively. Secondary structures are labeled as α1–α14 for helices and β1–β11 for strands. (B) Structure-based sequence alignment of McsB. Species abbreviations are as follows: Bs, *Bacillus subtilis*; Sa, *Staphylococcus aureus*; Gs, *Geobacillus*

*stearothermophilus*; Lm, *Listeria monocytogenes*; Bc, *Bacillus cereus*; Sc, *Shouchella clausii*; Hh, *Halalkalibacterium halodurans*. Blue triangles indicate the positions of Arg residues phosphorylated in *B. subtilis* McsB.

### Figure S7

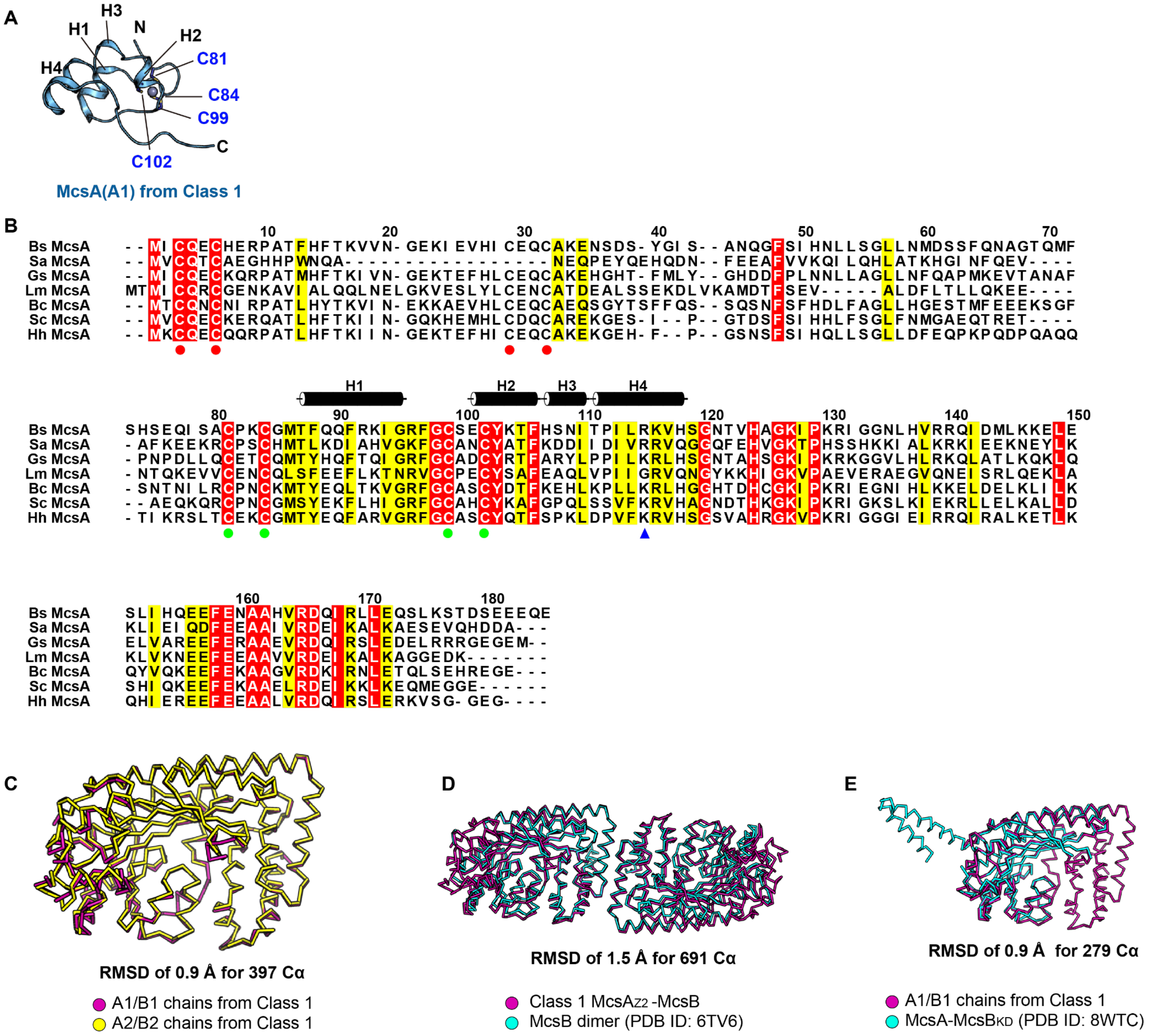


**Supplementary Figure 7. Structure of McsA$_{Z2}$ in the McsA$_{Z2}$-McsB complex.** (A) Ribbon diagram of chain A1 of McsA from the Class 1 cryo-EM structure. Helices are labeled H1–H4. Cys residues that coordinate with a zinc atom are labeled as C81, C84, C99, and C102. (B) Sequence alignments of McsA. Species abbreviations follow those shown in Fig. S6. Green dots indicate zinc-coordinating Cys residues. Blue triangle marks the position of the phosphorylated Arg residue in *B. subtilis* McsA. (C) Structural comparison of two McsA$_{Z2}$-McsB heterodimers in the Class 1 cryo-EM structure. (D) Structural comparison of the Class 1 structure with the crystal structure of the McsB dimer (PDB ID: 6TV6). (E) Structural comparison of McsA$_{Z2}$-McsB heterodimer (chains A1 and B1) from the Class 1 structure with the crystal structure of McsA-McsB$_{KD}$ (PDB ID: 8WTC).

**Figure S8**

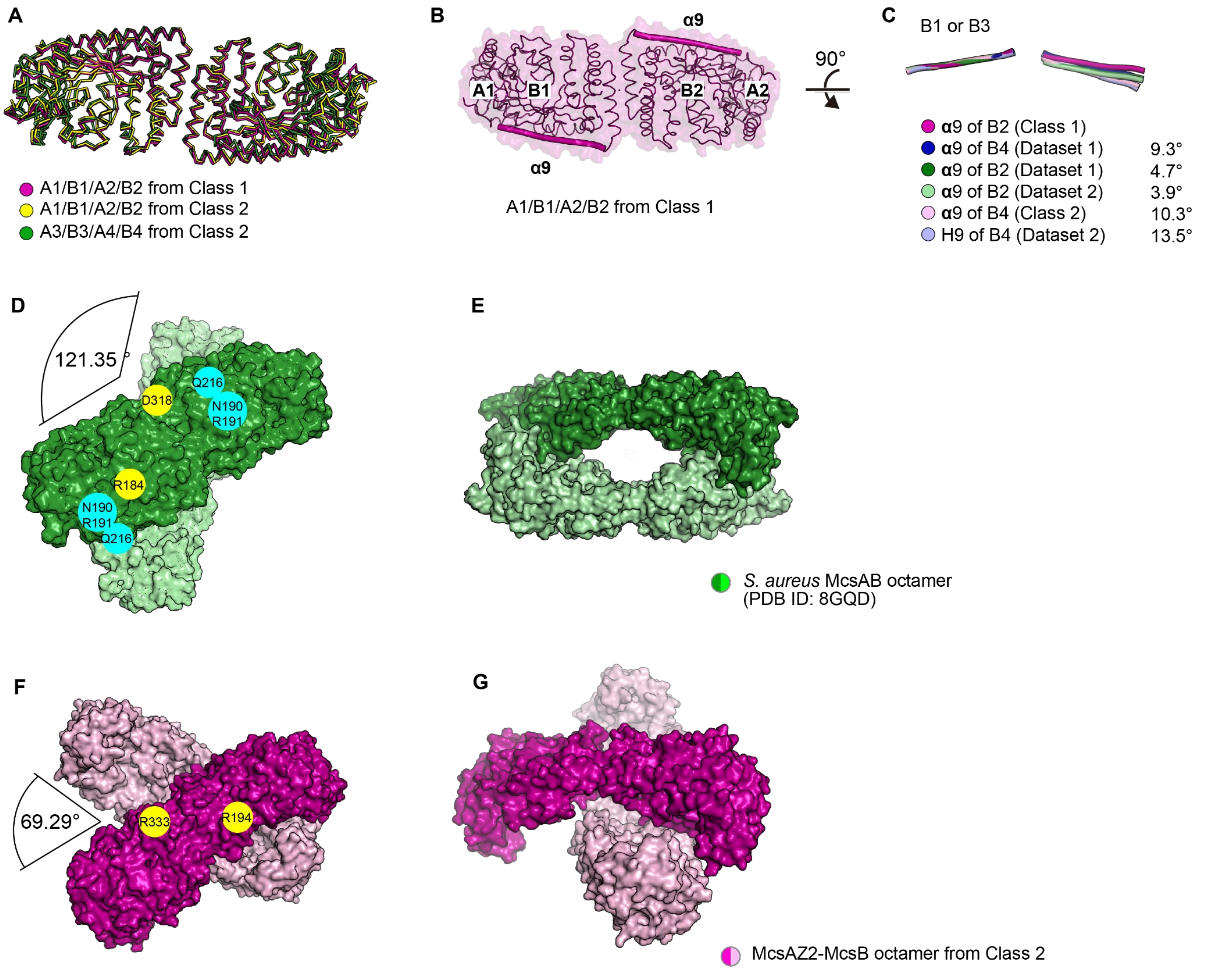


**Supplementary Figure 8. Class 2 cryo-EM structure of McsA$_{Z2}$-McsB.** (A) Structural comparison of three McsA$_{Z2}$-McsB heterotetramers from Class 1 and Class 2 structures. Root-mean-square deviation (RMSD) value between two heterotetramers of Class 2 is 1.9 Å for 798 Cα atoms. RMSD values between the Class 1 tetramer and the two Class 2 tetramers are 1.2 Å for 800 Cα atoms and 2.5 Å for 796 Cα atoms, respectively. (B, C) Conformational comparison of McsA$_{Z2}$-McsB heterotetramers from Class 1, Class 2, Dataset 1, and Dataset 2 structures. After superimposing one McsB subunit of the McsA$_{Z2}$-McsB heterotetramers, conformational differences among the other McsB subunits were compared by analyzing angle differences of the long helix α9. (D, E) Cryo-EM structure of *S. aureus* McsAB (PDB ID: 8GQD), which forms a symmetric dimer of the McsAB heterotetramer. (F, G) Class 2 cryo-EM structure of *B. subtilis* McsA$_{Z2}$-McsB. The magenta heterotetramer is shown in the same orientation as the dark green heterotetramer in panel D.

**Figure S9**

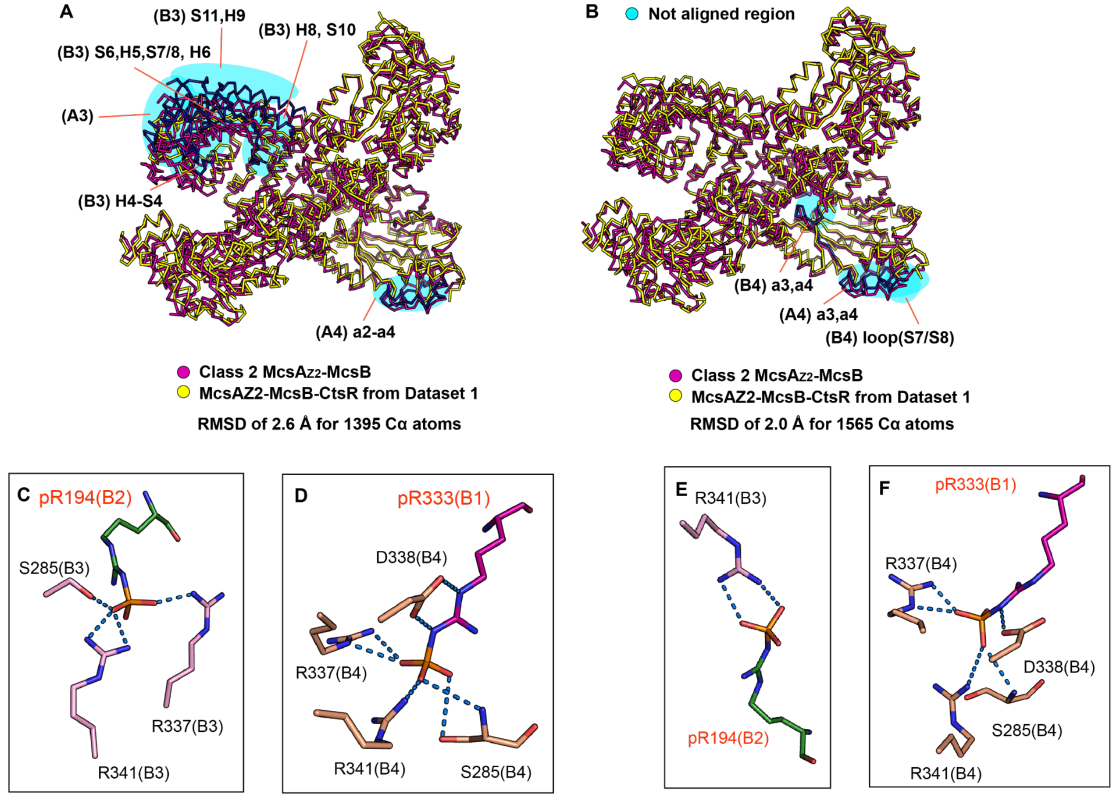


**Supplementary Figure 9. Cryo-EM structures of the McsA$_{Z2}$-McsB-CtsR complex**. (A) Structural comparison of McsA$_{Z2}$-McsB octamers from the Class 2 structure of McsA$_{Z2}$-McsB and the Dataset 1 structure of McsA$_{Z2}$-McsB-CtsR complex. Regions that are not structurally aligned are highlighted in cyan. (B) Structural comparison of McsA$_{Z2}$-McsB octamers from the Class 2 structure of McsA$_{Z2}$-McsB and the Dataset 2 structure of McsA$_{Z2}$-McsB-CtsR complex. (C, D) Stick models illustrating interactions between the pArg and pocket of McsB in Dataset 1 structure: pR194 (C) and pR333 (D). (E, F) Stick models illustrating interactions between the pArg and pocket of McsB in Dataset 2 structure: pR194 (E) and pR333 (F).

**Figure S10**

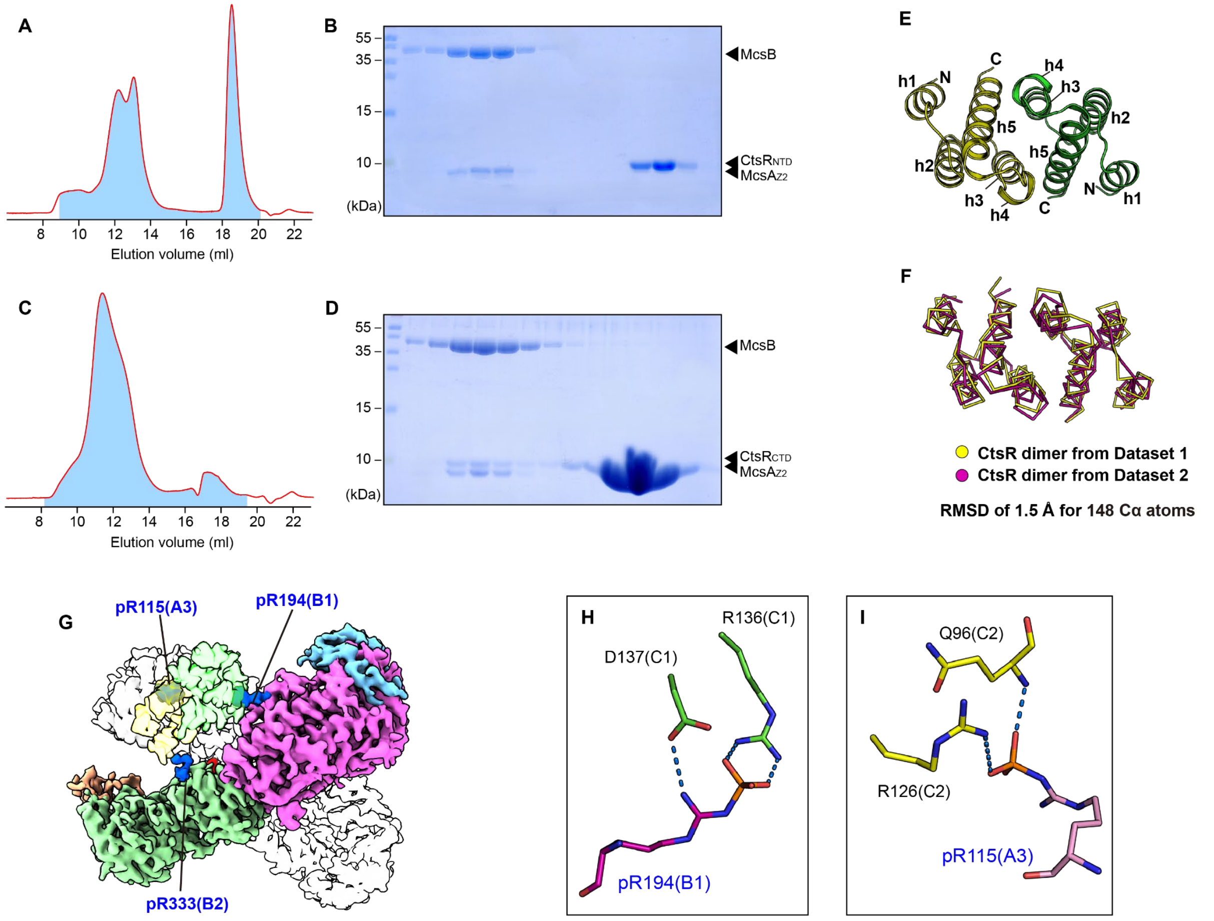


**Supplementary Figure 10. Interactions between McsA$_{Z2}$-McsB and CtsR.** (A, B) Absorbance curve at 280 nm (A) and SDS-PAGE analysis (B) showing the size-exclusion chromatography (SEC) results of McsA$_{Z2}$-McsB with excess CtsR$_{NTD}$. The blue-shaded area under the absorbance curve corresponds to the SEC fractions analyzed using SDS-PAGE. (C, D) Absorbance curve at 280 nm (C) and SDS-PAGE analysis (D) showing the SEC results of McsA$_{Z2}$-McsB with excess CtsR$_{CTD}$. (E) Ribbon diagram showing the structure of the CtsR dimer from Dataset 1. Helices are labeled h1–h5. (F) Structural comparison of CtsR dimers from Dataset 1 and Dataset 2 structures. (G) Dataset 2 cryo-EM density map showing the position of three pArg residues for CtsR binding at different orientations. (H, I) Stick models illustrating interactions between the pArg and CtsR in Dataset 1 cryo-EM structure: pR194 of McsB (H) and pR115 of McsA (I).

Figure S11

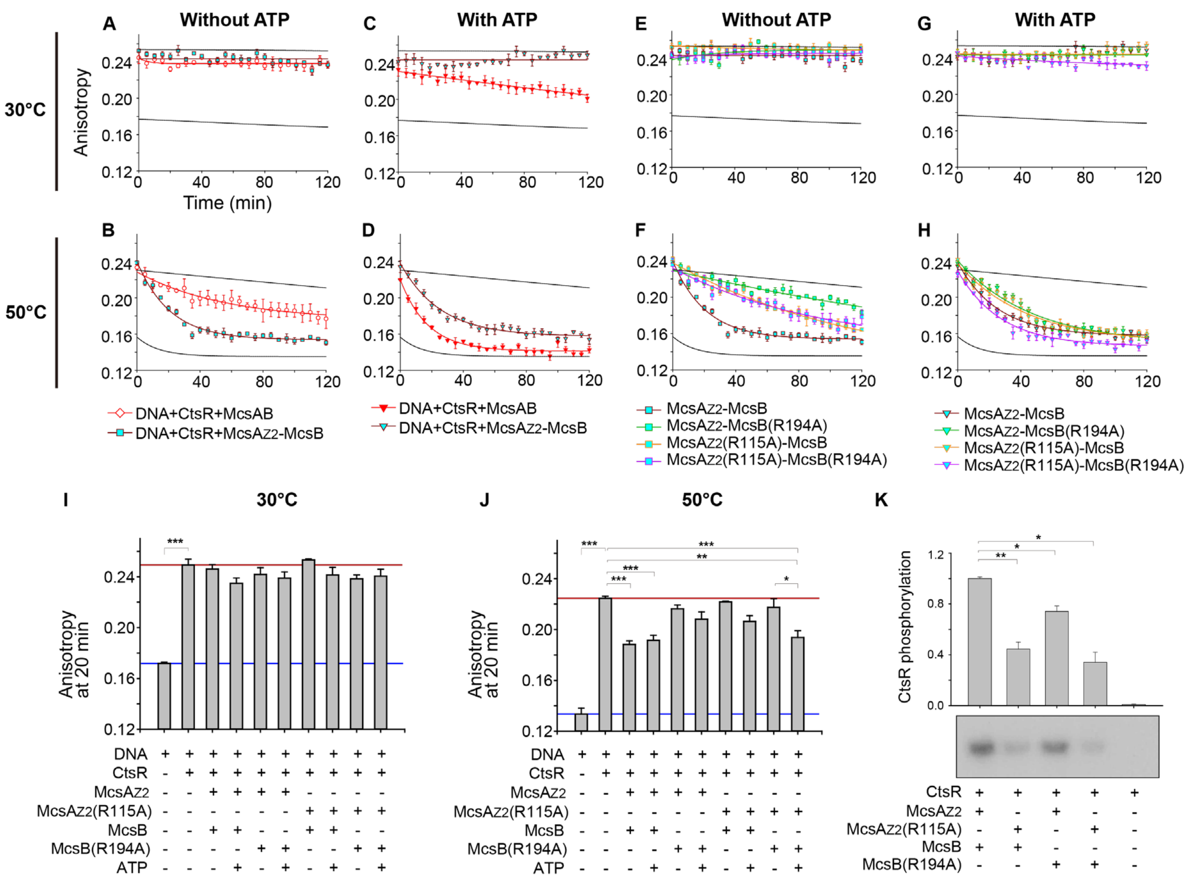


**Supplementary Figure 12. Autophosphorylation-dependent hijacking of CtsR by McsA$_{Z2}$-McsB during heat shock.** (A, B) Anisotropy measurements of the CtsR-operator complex in the presence of either McsAB or McsA$_{Z2}$-McsB. The anisotropy was measured without ATP for 2 h at 30 °C (A) and 50 °C (B). (C, D) Anisotropy measurements of the CtsR-operator complex in the presence of McsAB or McsA$_{Z2}$-McsB. Anisotropy was measured with ATP for 2 h at 30 °C (C) and 50 °C (D). (E, F) Anisotropy measurements of the CtsR-operator complex in the presence of McsA$_{Z2}$-McsB or its mutants. Anisotropy was measured without ATP for 2 h at 30 °C (E) and 50 °C (F). (G, H) Anisotropy measurements of the CtsR-operator complex in the presence of McsA$_{Z2}$-McsB or its mutants. Anisotropy was measured with ATP for 2 h at 30 °C (G) and 50 °C (H). (I, J) Bar graphs showing anisotropy values at 20 min at 30 °C (I) and 50 °C (J). Anisotropy data are presented as the mean ± standard error from three independent experiments, each with three replicates. (K) Kinase activity of McsA$_{Z2}$-McsB and its mutants, assessed by measuring CtsR phosphorylation in the presence of McsA$_{Z2}$-McsB or its mutants. Data are presented as the mean ± standard error from three independent experiments. Asterisks in panels (I–K) indicate statistical significance levels: *$p < 0.05$, **$p < 0.01$, and ***$p < 0.001$.

**Figure S12**

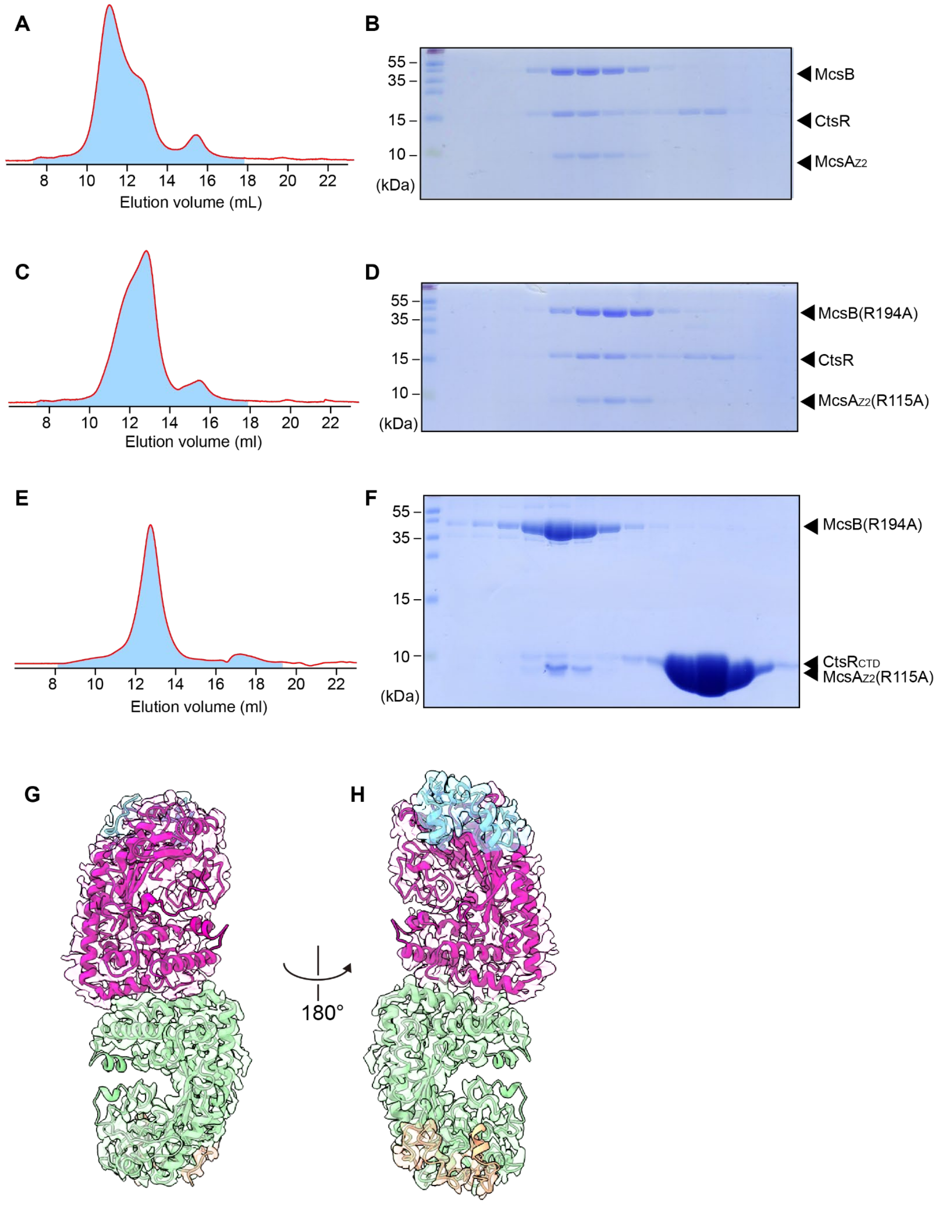


**Supplementary Figure 12. Interactions between McsA$_{Z2}$-McsB mutants and CtsR.** (A, B) Absorbance curve at 280 nm (A) and SDS-PAGE analysis (B) showing the SEC results of McsA$_{Z2}$-McsB with excess CtsR. (C, D) Absorbance curve at 280 nm (C) and SDS-PAGE analysis (D) showing the SEC result of McsA$_{Z2}$(R115A)-McsB(R194A) with excess CtsR. (E, F) Absorbance curve at 280 nm (E) and SDS-PAGE (F) showing the SEC result of the McsA$_{Z2}$(R115A)-McsB(R194A) complex with excess CtsR$_{CTD}$. (G, H) Cryo-EM density map of the McsA$_{Z2}$(R115A)-McsB(R194A)-CtsR complex superimposed with the McsA$_{Z2}$-McsB heterotetramer.

**Supplementary Table 1. Data-collection and refinement statistics for the structure determination**

| Data set | $McsA_{Z2}$-McsB | | $McsA_{Z2}$-McsB-CtsR Set 1 | $McsA_{Z2}$-McsB-CtsR Set 2 |
|---|---|---|---|---|
| **Data collection** | | | | |
| Microscope | Titan Krios G4 | | Titan Krios G4 | Titan Krios G4 |
| Detector | Falcon4 | | Falcon4 | Falcon4 |
| Magnification | 165,000 | | 165,000 | 165,000 |
| Voltage (kV) | 300 | | 300 | 300 |
| Electron Exposure ($e^-/Å^2$) | 50 | | 50 | 50 |
| Defocus Range (μm) | -0.6 to -2.0 | | -0.6 to -2.0 | -0.6 to -2.0 |
| Pixel size (Å) | 0.7451 | | 0.7451 | 0.7451 |
| No. frames/movie | 30 | | 60 | 36 |
| Total exposure time (sec) | 5.24 | | 6.35 | 5.23 |
| Number of movies | 5,903 | | 2,498 | 12,050 |
| **3D density map reconstruction** | Class I | Class II | | |
| Symmetry imposed | *C*1 | *C*1 | *C*1 | *C*1 |
| No. initial particles | 3,724,487 | 3,724,487 | 1,966,040 | 7,576,165 |
| No. final particles | 228,764 | 266,396 | 15,254 | 36,545 |
| FSC threshold | 0.143 | 0.143 | 0.143 | 0.143 |
| Map resolution (Å) | 2.89 | 3.02 | 3.56 | 3.38 |
| **Model refinement** | Class I | Class II | | |
| Composition | | | | |
| Atoms | 6,360 | 12,788 | 14,155 | 14,022 |
| Residues | 803 | 1,610 | 1,782 | 1,766 |
| Ligands | Zn: 2 | Zn: 4 | Zn: 4 | Zn: 4 |
| Bonds (RMSD) | | | | |
| Length (Å) | 0.019 | 0.004 | 0.009 | 0.007 |
| Angle (°) | 1.299 | 0.568 | 0.958 | 0.868 |
| Mean B-factors | | | | |
| Protein | 45.97 | 40.81 | 81.76 | 98.65 |
| Ligand | 50.60 | 52.64 | 101.36 | 127.63 |
| Ramachandran plot (%) | | | | |
| Favored | 96.48 | 93.62 | 94.74 | 94.80 |
| Allowed | 3.52 | 6.38 | 5.26 | 5.14 |
| Outliers | 0.00 | 0.00 | 0.00 | 0.06 |
| Rotamer outliers (%) | 4.82 | 0.35 | 0.25 | 0.58 |
| Cβ outliers (%) | 0.00 | 0.00 | 0.00 | 0.00 |
| Clash score | 6.03 | 8.54 | 19.37 | 15.87 |
| MolProbity score | 2.08 | 1.88 | 2.15 | 2.06 |
| **Model vs. Data** | | | | |
| CC (mask) | 0.81 | 0.77 | 0.76 | 0.82 |
| CC (box) | 0.66 | 0.68 | 0.72 | 0.74 |
| CC (peaks) | 0.64 | 0.64 | 0.63 | 0.67 |
| CC (volume) | 0.76 | 0.73 | 0.74 | 0.80 |
| CC for ligands | 0.35 | 0.33 | 0.47 | 0.58 |
